# Modelling collagen fibril self-assembly from extracellular medium in embryonic tendon

**DOI:** 10.1101/2023.03.13.532430

**Authors:** Christopher K. Revell, Jeremy A. Herrera, Craig Lawless, Yinhui Lu, Karl E. Kadler, Joan Chang, Oliver E. Jensen

## Abstract

Collagen is a key structural component of multicellular organisms and is arranged in a highly organised manner. In structural tissues such as tendons, collagen forms bundles of parallel fibres between cells, which appear within a 24 hour window between E13.5 and E14.5 during mouse embryonic development. Current models assume that the organised structure of collagen requires direct cellular control, whereby cells actively lay down collagen fibrils from cell surfaces. However, such models appear incompatible with the time- and length-scales of fibril formation. We propose a phase-transition model to account for the rapid development of ordered fibrils in embryonic tendon, reducing reliance on active cellular processes. We develop phase-field crystal simulations of collagen fibrillogenesis in domains derived from electron micrographs of inter-cellular spaces in embryonic tendon and compare results qualitatively and quantitatively to observed patterns of fibril formation. To test the prediction of this phase-transition model that free protomeric collagen should exist in the intercellular spaces prior to the formation of observable fibrils, we use laser-capture microdissection, coupled with mass spectrometry, which demonstrates steadily increasing free collagen in intercellular spaces up to E13.5, followed by a rapid reduction of free collagen that coincides with the appearance of less soluble collagen fibrils. The model and measurements together provide evidence for extracellular self-assembly of collagen fibrils in embryonic mouse tendon, supporting an additional mechanism for rapid collagen fibril formation during embryonic development.

## 1 Introduction

Collagen is the largest and most abundant protein found in vertebrates [1] and forms the primary structural component of multicellular tissues across multiple scales, ranging from the cornea to tendon and skin. Despite the critical importance of collagen to multicellular life, and its role in conditions such as Ehlers–Danlos syndrome [2] and osteogenesis imperfecta [3], there remain significant gaps in our understanding of the initial development of collagen fibrils [4], especially in the mesoscale regime at scales above that of chemical bonds but below that of organs and tissues.

Previous cross-section electron microscope images of embryonic mouse tail tendon suggest that fibrils appear within a 24-hour window between embryonic day 13.5 and 14.5 (E13.5-14.5), forming parallel fibres that appear in cross section as a roughly hexagonal lattice of approximately identical circles [Figure 1]. This lattice develops in inter-cellular spaces, which are themselves extended tubes parallel to the long axis of the tendon [5]. The highly ordered nature of fibrils within tendon led to the assumption that cells must exert tight active control over their development, and thus fibrils were hypothesised to be actively laid down only at cell surfaces [6]. This requires that cells retain contact with the collagen fibril and grow it in multiple ways: 1) by continuously feeding new collagen protomer at the contact site, which would require the fibrils to move extensively through the inter-cellular space as the fibril grows [7]; 2) cells may instead move along the fibril with the growing tip embedded within the cell; or 3) cells remain *in situ* with several smaller fibrils assembled at cell surfaces which are subsequently attached together end-to-end [8]. The cell-mediated fibrillogenesis hypothesis is supported by observations of structures known as fibripositors, within which collagen fibrils were shown to be embedded within a cell [9]. However, the existence of fibripositors was previously only observed at E14.5 and not E13.5, when fibrils have already extensively occupied the intercellular spaces [9]. Further, highly ordered collagen fibrils observed at the centre of dense fibril bundles [Figure 1b], where cell-surfaces are not in close proximity, also raises the question of how cells may coordinate rapid but precise movements (either the cell body or multiple fibril ends) that transform the intercellular spaces from voids to dense ordered bundles of collagen fibrils, all within 24 hours. Here, we explore a mechanism complementing cell-mediated fibrillogenesis, by considering the rapid appearance of ordered fibril arrays within a disordered medium as a phase transition or crystallisation process [10].

**Figure 1:**
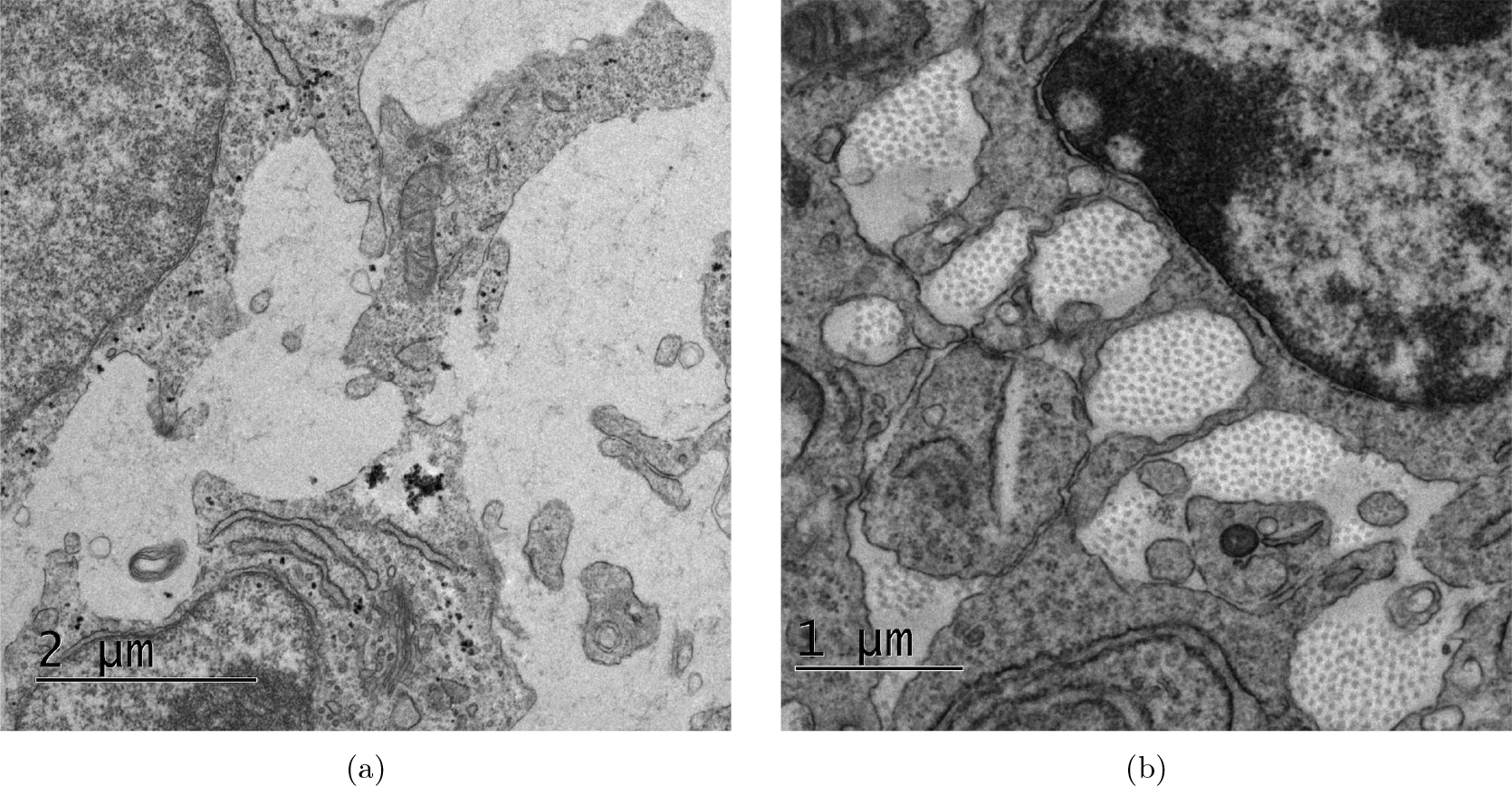
(a) Cross section of mouse embryonic tail tendon at E13.5, showing absence of fibrils in inter-cellular spaces. (b) Cross section of mouse embryonic tail tendon at E14.5, demonstrating sudden appearance of fibril lattices in inter-cellular spaces 24 hours later. Images are unpublished from study [9].

We investigate this hypothesis with a combined experimental and theoretical approach. We simulate self-assembly of fibrils using a phase-field crystal (PFC) model, a standard mathematical description of self-assembly, as outlined in Section 2.1. We perform simulations in irregular domains derived from intercellular spaces observed in EM images of mouse tail tendon at the stage of development when fibrils first appear. We compare the results of these simulations to corresponding patterns of fibrils observed directly in these EM images, and discuss similarities and differences, in particular noting the density of defects and the presence of voids in fibril packing.

Self-assembly of collagen fibrils is predicated upon the hypothesis that protomeric collagen is secreted by cells into the inter-cellular space prior to the formation of visible collagen fibrils. Crystallisation of this extracellular pool of collagen protomers is in contrast to previous suggestions of active assembly of fibrils by cells at cell surfaces [6], but does not preclude the involvement of cell-directed fibrillogenesis in this process. Rather, it may work in tandem to aid rapid growth of fibrils during embryonic development. We validate this hypothesis by experimental observation of the contents of inter-cellular spaces in embryonic tendon, using laser-capture microdissection coupled with mass spectrometry analysis (LCM-MS) [11].

The results suggest a steady increase of collagen and proteins involved in collagen fibrillogenesis over time up to E13.5, followed by a sharp reduction of available collagen in the inter-cellular space between the time points where collagen fibrils appear in the tendon tissue. This supports the phase-transition model proposed here, where the available collagen, easy to extract for LCM-MS [12], peaks prior to the formation of observable fibrils, when protomers are incorporated into less soluble collagen fibrils in a crystallisation process.

## 2 Materials and Methods

### 2.1 Mathematical modelling

#### The phase-field crystal model

We hypothesise that fibrils form by crystallisation of an existing pool of collagen protomers secreted into the inter-cellular space prior to the visible appearance of fibrils, with fibril ordering determined by the geometry of the inter-cellular space. We model the extracellular self-assembly of collagen protomers into fibrils using the phase field crystal (PFC) equation [13, 14, 15] [Appendix A], a sixth-order nonlinear partial differential equation that has been used extensively to model the emergence of ordered structures from a disordered medium [16], including crystal nucleation in colloidal suspensions [17, 18] and even the internal structure of collagen fibrils themselves [19, 20]. The PFC equation is a coarse-grained model derived from explicit inter-particle interactions [21, 22]; this coarse-graining ensures that the PFC method is scalable to larger systems while remaining grounded in nano-scale physics. However, rather than formally derive a free energy specific to collagen from first principles, we assume a canonical functional form with a minimal number of parameters. To reduce computational cost, our simulations describe evolution in a two-dimensional plane normal to a fibril bundle, neglecting variation along the long axis of inter-cellular spaces. In addition to symmetry along the long axis of inter-cellular spaces [5], we assume that the transverse isotropy of resulting fibril arrays is enhanced by a variety of factors, such as nematic ordering of free collagen protomers [23], geometric confinement [24], or longitudinal mechanical stress [25, 26, 27] that together promote nematic alignment of fibrils along the axis of a developing tendon. Explicitly modelling 3D alignment using anisotropic forms of the PFC [28, 29, 30] was considered to be beyond the scope of this study.

#### The phase field

A phase field is a function that divides a domain into two primary states, labelled with different scalar values, in this case crystallised collagen fibril (in which collagen is highly condensed) and extracellular medium (in which collagen protomers exist at relatively low concentration). More precisely, the PFC model describes the evolution of a spatial phase field *ϕ* (***x***, *t*), where *ϕ* varies in the range −1 to 1. The evolving phase field divides the solution domain into regions Ω*_±_* for which *ϕ* (***x***, *t*) *≈ ±*1 for **x** *∈* Ω*_±_*, with narrow interfaces over which *ϕ* varies with a sharp gradient. We use *ϕ* (***x***, *t*) *≈* −1 to indicate that fibrillar collagen exists at position ***x*** *∈* Ω_−_ and time *t*, and *ϕ* (***x***, *t*) *≈* 1 to indicate that protomeric collagen suspended in extracellular medium exists at ***x*** *∈* Ω_+_. The PFC equation models the phase separation of these two states, and consequently the crystallisation of fibrillar collagen from an initial state. The phase field measures density fluctuations, so that the quantity 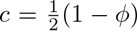 provides a crude (dimensionless) proxy for molecular concentration: we consider 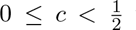 to define regions occupied by protomer (with low molecular density) and regions with 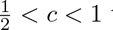 to define fibrillar regions in which collagen is densely aggregated. Phase separation is driven by gradients of free energy; however the system exhibits glassy dynamics, meaning that multiple metastable states can arise, depending on fine details of the initial conditions in any realisation of the model. Given the rapid diffusion of protomeric collagen [31] relative to the timescale of fibril assembly and lengthscale of the inter-cellular space, we assume a roughly uniform initial *ϕ* field with a small random fluctuation specified by a Gaussian random field. This disordered initial state leads us to take a statistical view of the fibril patterns that emerge.

#### Parameters

Spatial scales of the PFC are set by a parameter *q^∗^* that is adjusted to fit length scales measured via microscopy, as explained in Appendix A. Additionally, the PFC model has two dimensionless parameters (*r* and *ϕ*_0_) that can be related to the free energy that drives fibril assembly [Appendix A]. The parameter *r* measures the destabilising component of the free energy (relative to a stabilising component that penalises density gradients); *ϕ*_0_ measures the mean phase field over the domain (effectively specifying the initial collagen concentration). These parameters regulate the spatial patterns formed by steady-state solutions of the PFC model [13]. Within certain ranges, however, predicted fibril patterns are statistically robust with respect to parameter variations. The only additional parameters are a correlation length *λ* and variance *m*^2^ of the initial state, specified as a Gaussian random field [32]; again, predicted patterns show little sensitivity to reasonable variations of these parameters.

#### Computational methods

We applied a split semi-implicit approach to solving the PFC equation [33], using the Julia differential equations library [34, 35] to achieve good computational performance [Appendix B]. To accommodate potential fibril nucleation at domain boundaries, fibril self-assembly was modelled by solving the PFC equation in irregular domains defined by inter-cellular spaces, extracted from electron microscope (EM) images of embryonic mouse tail tendons in cross-section [Appendix C], allowing the influence of domain shape on fibril patterns to be investigated. To compare fibril patterns in EM images to fibril patterns in simulations, we mapped patterns of spatial defects by using a Delaunay triangulation to identify the number of nearest neighbours of each fibril. We measured deviations from the expected number of neighbours to quantify the irregularities in the packing and to analyse the distribution of fibril neighbour separation and neighbour counts.

We selected values for parameters *λ* and *m*^2^ based on initial sense-testing, confirming that these parameters did not significantly affect equilibrium states of the PFC equation, selected a time step *dt* that ensured stability, and ran to a maximum time that ensured a near-equilibrium final state [Appendix D]. We then ran many simulations in a wide region of (*r, ϕ*_0_)-space to investigate the set of possible outcomes from random initial conditions.

### 2.2 Image analysis

We applied a similar image analysis pipeline to both EM images and simulation results [Appendix E]. Starting from a representative set of EM images showing inter-cellular spaces and collagen fibril bundles, we developed a pipeline to manually identify locations of fibrils. Manual selection was preferred over segmentation due to limitations of noisy EM images. We performed a Delaunay triangulation [36] over this set of points for each image [Appendix E]. With these Delaunay triangulations, we were able to analyse the patterns of nearest neighbours for each image, including the distribution of spacing between nearest neighbours, and the distribution of neighbour counts for each fibril.

The neighbour count of each fibril within the triangulation reveals defects in the lattice pattern [37]. We visualised the neighbour counts of fibrils as a Voronoi tessellation [36], with each Voronoi cell representing one fibril, and the colour of the cell being clear if the fibril has 6 neighbours, red if the fibril has 5 neighbours or blue if the fibril has 7 neighbours. We calculated the defect proportion (*d*) for each image by taking the ratio of the number of fibrils that do not have 6 neighbours to the total number of fibrils, excluding those on the periphery of the bundle.

### 2.3 Biological samples

#### Mice

The care and use of all mice in this study were carried out in accordance with the UK Home Office Regulations, UK Animals (Scientific Procedures) Act of 1986 under the Home Office License (PP3720525). 12-week old female C57BL/6 mice were time-mated with males that were removed after overnight housing. Embryo ages were calculated as day 0 on the morning where females were plugged, and embryos were isolated at ages 12.5 days, 13 days, 13.5 days, 14 days, 14.5 days (E12.5, E13, E13.5, E14, E14.5). Embryos were randomly sampled for each analysis technique. Embryos were kept in ice-cold PBS and tail tendons were removed at the base. These tail tendons were then fixed with 4 % formalin for laser-capture micro-dissection, or electron microscopy (EM) fixative (2 % glutaraldehyde/100mM phosphate buffer at pH 7.2).

#### Electron microscopy

After fixation, the tails were washed in ddH_2_O for 5 minutes, repeated 3 times. The samples were then transferred to 2 % osmium (v/v)/1.5 % potassium ferrocyanide (w/v) in cacodylate buffer (100 mM, pH 7.2) and further fixed for 1 h, followed by extensive washing in ddH_2_O. This was followed by 40 minutes of incubation in 1 % tannic acid (w/v) in 100 mM cacodylate buffer, and then extensive washes in ddH_2_O. Samples were then placed in 2 % osmium tetroxide for 40 min, followed by extensive washes in ddH_2_O. Samples were incubated with 1 % uranyl acetate (aqueous) at 4 *^◦^*C for at least 16 h, and then washed again in ddH_2_O. Samples were then dehydrated in graded ethanol in the following regime: 30 %, 50 %, 70 %, 90 % (all v/v in ddH_2_O) for 8 min at each step. Samples were then washed 4 times in 100 % ethanol, and transferred to pure acetone for 10 min. The samples were then infiltrated in graded series of Agar100Hard in acetone (all v/v) in the following regime: 30 % for 1 h, 50 % for 1 h, 75 % for 16 h, 100 % for 5 h. Samples were then transferred to fresh 100 % Agar100Hard in labeled moulds and allowed to cure at 60 *^◦^*C for 72 h. Sections (80 nm) were cut and examined using a Tecnai 12 BioTwin electron microscope.

#### Histological staining and imaging

5 *µ*m sections of formalin-fixed and paraffin-embedded (FFPE) mouse tail specimens were H&E-stained by using an automated stainer (Leica XL) at University of Manchester’s Histology Core as previously described [11, 38]. We used a DMC2900 Leica instrument with Leica Application Suite X software for imaging.

#### Laser-capture microdissection (LCM) of mouse tail tendons

A mouse tail contains 4 tendon bundles that are uniformly spaced around a central bone [39] [Figure 4a]. The MMI CellCut Laser Microdissection System (Molecular Machines & Industries) was used in combination with their MMI CapLift technology to capture regions of interest within embryonic tail tendon bundles on MMI membrane slides (MMI, 50102) as previously described [11, 38]. Laser power was set to be between 30-40 %, cut speed at 50 *µ*m/sec and z drill at 5 *µ*m. In Figure 4b we show our LCM capacity to cut and capture the 4 tendons at embryonic timepoints E12.5 and E14.5. In this study, we capture tail tendons for five timepoints E12.5, E13.0, E13.5, E14.0 and E14.5 (n=3 mice per timepoint; a total of 15 samples) with a tissue collection volume of roughly 0.01 mm^3^ per sample. Samples were stored in 4 *^◦^*C prior to laser-capture and stored at −80 *^◦^*C once tendons were collected. The tendons were subject to mass spectrometry preparation. In short, samples underwent a multistep process to maximize protein yield, including high detergent treatment, heating and physical disruption, as previously described [11, 38].

#### Liquid chromatography coupled tandem mass spectrometry

The separation was performed on a Thermo RSLC system (ThermoFisher), as previously described [40]. The analytical column was connected to a Thermo Exploris 480 mass spectrometry system via a Thermo nanospray Flex Ion source via a 20 µm ID fused silica capillary. The capillary was connected to a fused silica spray tip with an outer diameter of 360 µm, an inner diameter of 20 µm, a tip orifice of 10 µm and a length of 63.5 mm (New Objective Silica Tip FS360-20-10-N-20-6.35CT) via a butt-to-butt connection in a steel union using a custom-made gold frit (Agar Scientific AGG2440A) to provide the electrical connection. The nanospray voltage was set at 1900 V and the ion transfer tube temperature set to 275 *^◦^*C.

Data were acquired in a data-dependent manner using a fixed cycle time of 1.5 s, an expected peak width of 15 s and a default charge state of 2. Full MS data was acquired in positive mode over a scan range of 300 to 1750m/z, with a resolution of 1.2 *×* 10^5^ FWHM, a normalized AGC target of 300 %, and a max fill time of 25 ms for a single microscan. Fragmentation data was obtained from signals with a charge state of +2 or +3 and an intensity over 5000 ion/second, and they were dynamically excluded from further analysis for a period of 15 s after a single acquisition within a 10 ppm window. Fragmentation spectra were acquired with a resolution of 1.5 *×* 10^4^ FWHM with a normalized collision energy of 30 %, a normalized AGC target of 300 %, first mass of 110m/z and a max fill time of 25 ms for a single microscan. All data were collected in profile mode.

## 3 Results

### Simulated fibril pattern formation and maturation

The short timescale [Figure 2a] and long timescale [Figure 2b] evolution of the PFC model shows rapid emergence of ordered structures followed by slow coarsening. From a random initial condition (mimicking release of collagen protomer from surrounding cells prior to the start of the simulation at *t* = 0), the pattern initiates at the boundary of the extracellular domain, propagating inwards towards its center [Figure 2a]. Having filled the domain (in this example), the pattern adjusts slowly over time, with the density of defects in the fibril packing falling slowly as the pattern takes on an increasingly crystalline structure [Figure 2b,d]. However, because of the irregularity of the boundary, and because some features of the initial random field are effectively frozen into the pattern, defects are a persistent long-term feature of the fibril array, and are present at a sufficiently high density to consider the pattern disordered. Defects appear as disclinations (isolated fibrils with 5 neighbours, coloured red, or with 7 neighbours, coloured blue in Figure 2a,b), as dislocations (red/blue pairs), and sometimes as linear pleats (or scars) containing an even (or odd) number of red/blue fibrils, which can be considered as grain boundaries [41]. Defects are not plotted for fibrils at the periphery of the cluster, leading to fluctuations in the defect proportion among interior fibrils in Figure 2d, which otherwise falls over time. The free energy driving the evolution falls monotonically as the system reaches a near-equilibrium state [Figure 2c]. Likewise, the availability of collagen protomer falls over time, as protomer is incorporated into fibrils [Figure 2c]. Here we have used 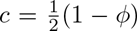 as proxy for collagen density, taking 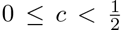 as representative of low-density regions (with collagen in protomeric form, red regions of the colour map in Figure 2a,b,e) and 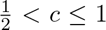 as representative of high density regions in which collagen aggregates into fibrils (blue regions of colour map). The integral of *c* over regions for which 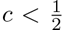 provides an estimate of the relative availability of protomeric collagen in the extracellular space.

**Figure 2:**
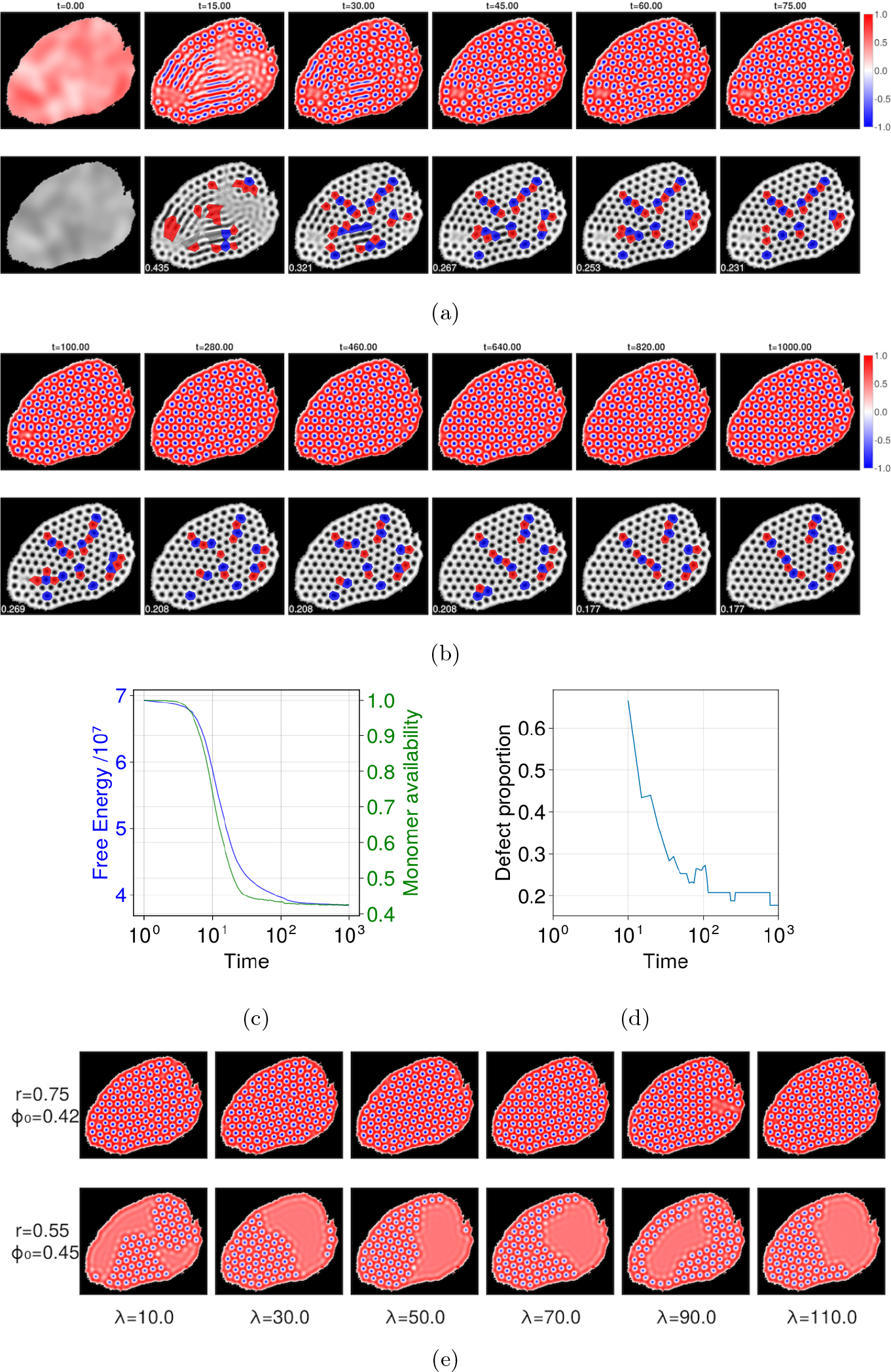
Time evolution of the phase field *ϕ*(**x**, *t*). (a) The short timescale dynamics shows rapid emergence of fibrillar structures, with corresponding defect analysis. The top row shows simulation data at 6 time points, with the colour showing the value of *ϕ*; the lower row shows the same data following a defect analysis as described in Section 2.2. A Voronoi tessellation is overlaid upon the fibril lattice, with clear cells for fibrils that have 6 neighbours, red cells for fibrils that have 5 neighbours, and blue cells for fibrils that have 7 neighbours (any other number of neighbours is shown with a grey cell). Each image is 881 nm in the horizontal dimension. (b) The long timescale dynamics shows slow coarsening of ordered structures towards the lowest accessible free-energy state. (c) Free energy and protomer availability (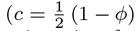, integrated over Ω_+_) against time for the system shown in panels a-b. (d) Defect proportion against time for the system shown in panels a-b. All data for a-d were generated with *r* = 0.8, *ϕ*_0_ = 0.4 and *λ* = 10. (e) Example simulation results showing the impact of varying the correlation length *λ* of the initial random field, for two sets of (*r, ϕ*_0_) values at *t* = 1000.

Simulation results are here presented with respect to dimensionless time units, time having been scaled relative to a factor involving free energy density, collagen mobility and a lengthscale. As the free energy is unknown, the model cannot make *a priori* predictions of the time needed for patterns to initially form. However it does predict that pattern maturation (involving gradual elimination of some defects) takes place over substantially longer timescales (ca. 500 time units) in comparison to the initial pattern formation (ca. 50 time units).

### Localised states and pattern robustness

Figure 2e illustrates the impact of varying the correlation length *λ* of the initial Gaussian random field. For fixed parameters (*r, ϕ*_0_), while there is variability between simulations as a result of the random initial conditions, *λ* does not appear to have a significant effect (at a statistical level) on large-time configurations. However the figure illustrates how voids can appear in the fibril array, a feature also evident in Figure 6 [Appendix D], which illustrates model predictions for a wider parameter sweep in *r* and *ϕ*_0_. Lower values of *ϕ*_0_ and higher values of *r* lead to inter-cellular spaces densely filled with fibril structures, while lower *r* and higher *ϕ*_0_ produces regions of dense structures with large gaps of empty space. These can be considered as so-called localised states, a characteristic feature of PFC models [13], whereby fibril patterns can co-exist with a homogeneous density field. As each simulation is run from a distinct random initial condition, the shape and location of voids in the pattern is highly variable, although the size of voids is regulated by parameter values. Correspondingly, there is variability in the density and distribution of defects [Figure 7, Appendix E], although these commonly appear in pairs (as dislocations) or lines (as scars or pleats). In summary, within the range of parameters investigated, the free-energy parameter *r* and the protomer abundance parameter *ϕ*_0_ primarily regulate the appearance of voids in the pattern, while stochastic effects determine the precise arrangements of defects (and voids) within the fibril arrays. Consequently, we use below *r* = 0.8 and *ϕ*_0_ = 0.4 as representative of parameter values leading to complete filling of the extracellular space with fibrils. In the model, only a single lengthscale parameter (*q^∗^*) was adjusted to fit model predictions to data.

### Defect analysis

To compare model predictions with observation, we identified the locations of fibrils within EM images of mouse tail tendon [Figure 3a], identified the patterns of defects [Figure 3b] and ran simulations in a domain matching that of the image [Figure 3c, d]. Although the stochastic nature of the simulation precludes recovery of precise details of the pattern, the configuration of defects is broadly similar. Simulations for the range of (*r, ϕ*_0_) values used in Figure 6 below generated predictions of the normalised distributions of nearest-neighbour separation [Figure 3g] and of fibril neighbour count [Figure 3h]. The predicted edge-length distribution is narrower than that measured in Figure 3a, b, indicating that the model underestimates the geometric disorder in the images, but the neighbour-count distribution is very similar.

**Figure 3:**
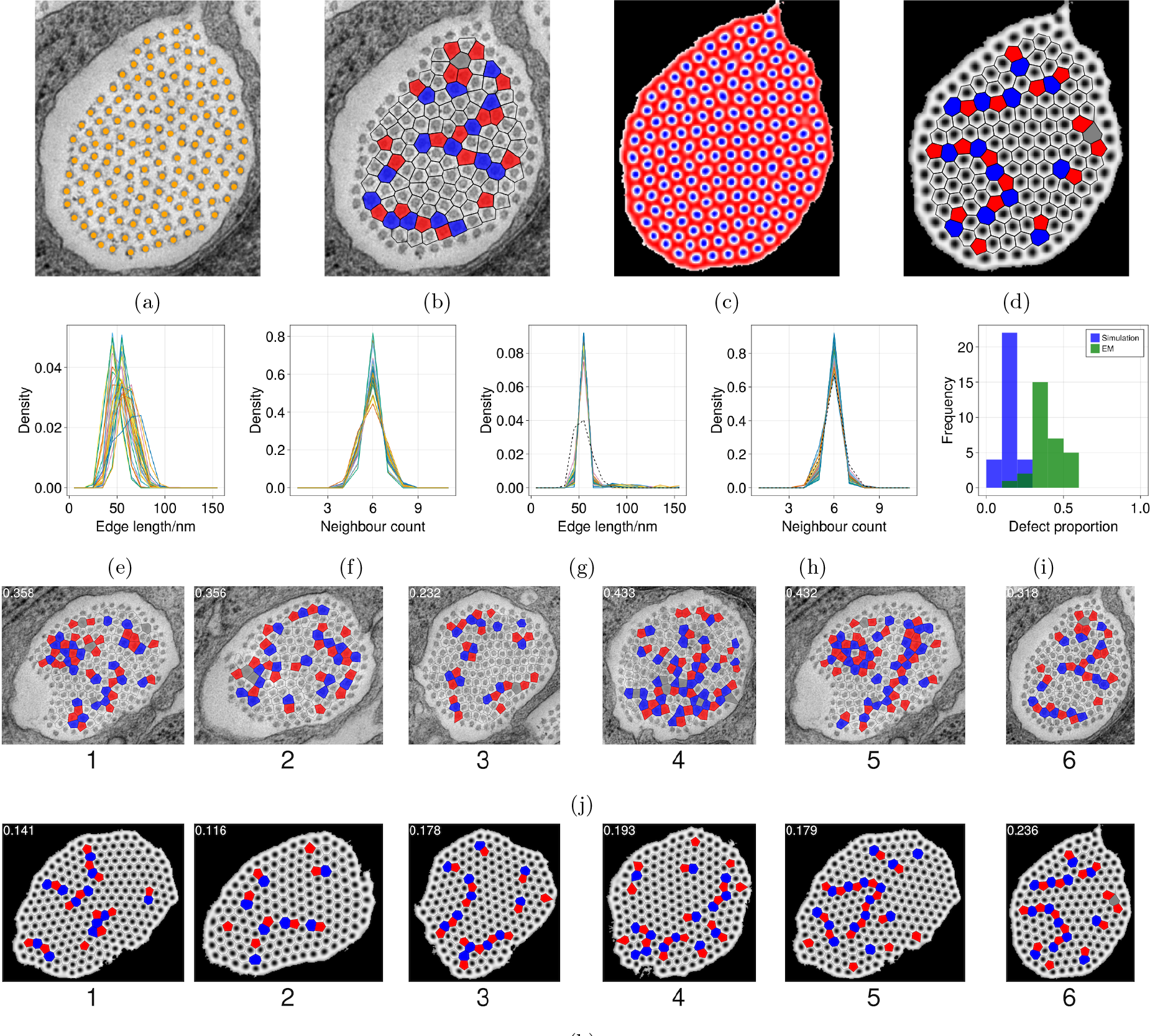
(a) Example of an EM image of an inter-cellular space with fibril locations highlighted by orange dots. (b) Corresponding image with defects in the fibril lattice highlighted such that fibrils with 5 neighbours are coloured red and those with 7 neighbours are coloured blue, excluding peripheral fibrils. (c) Final state of a simulation performed in a domain corresponding to the inter-cellular space in a, with parameters *r* = 0.8 and *ϕ*_0_ = 0.4. (d) Voronoi tessellation of result in c with Voronoi cells coloured by neighbour count as in b. (e) Normalised distributions of fibril nearest neighbour spatial separation for all 30 images shown in Figure 8. (f) Normalised distribution of the nearest neighbour counts for all internal fibrils, for all 30 images shown in Figure 8. (g) Set of normalised distributions of fibril nearest-neighbour-separation distance (edge length) for simulations at 36 points in *r* and *ϕ*_0_ space, run in a domain corresponding to inter-cellular space from image a, with length distribution observed in EM image shown with dotted line. (h) Set of normalised distributions of fibril nearest-neighbour count for simulations at 36 points in *r* and *ϕ*_0_ space, run in a domain corresponding to inter-cellular space from image a, with neighbour count distribution observed in EM image shown with dotted line. (i) Histogram of defect proportions across 30 tested EM images (green) and 30 simulations (blue) in corresponding inter-cellular spaces with parameters *r* = 0.8 and *ϕ* = 0.4. (j) A subset of EM images from Figure 8 with defects in fibril lattice visualised as in b. (k) A subset of simulation results from Figure 9 run in inter-cellular spaces corresponding to j with *r* = 0.8 and *ϕ* = 0.4, with defects in fibril lattice visualised as in d. All plots were generated using the Makie.jl plotting library [42].

To provide a wider comparison of model predictions against observation, 30 EM images showing inter-cellular spaces that contain collagen fibril bundles [Figure 8, Appendix F], of which 6 are shown in Figure 3j and 3k, were analysed to reveal patterns of defects and to recover normalised distributions of nearest-neighbour separation [Figure 3e], fibril neighbour count [Figure 3f] and defect density [Figure 3i]. Whilst there is some variation between images, there is a peak in median fibril spacing around 60nm, in line with past fibril diameter measurements [43, 44], and a sharp peak in neighbour count at 6 neighbours, as expected for a primarily hexagonal lattice. Simulations in the same 30 domains [Figure 9] predicted defect patterns arising at lower density [Figure 3i], indicating that the model fails to fully capture the topological disorder present in the EM images. In summary, simulations predict patterns of fibril arrangements that share many features of EM images, including localised states leading to voids in the fibril arrays, realistic defect patterns (albeit at a lower density on average than in images), and they provide evidence that rapid fibril formation (reflected by a rapid drop in available protomer) is likely to be followed by slower adjustment of patterns in which the defect density falls slightly.

### Collagens and other proteins essential for collagen synthesis increase over time prior to the formation of visible fibrils

To test the model prediction that protomeric collagen should exist in inter-cellular spaces prior to the formation of visible fibrils, we used mass spectrometry for protein quantification. We dissected tails from embryos at different stages of development, spanning the ages where collagen fibril formation occurs (E12.5-E14.5), and performed EM analyses to confirm the presence or absence of fibrils along the developmental time series. We observed the emergence of fibrils at E13.5. To identify proteins within regions of collagen fascicles, the major structural component of the tendon [Figure 4a], both before and after the appearance of collagen bundles, we utilised laser-capture microdissection to specifically isolate the fascicle regions for mass spectrometry analyses [Figure 4b].

**Figure 4:**
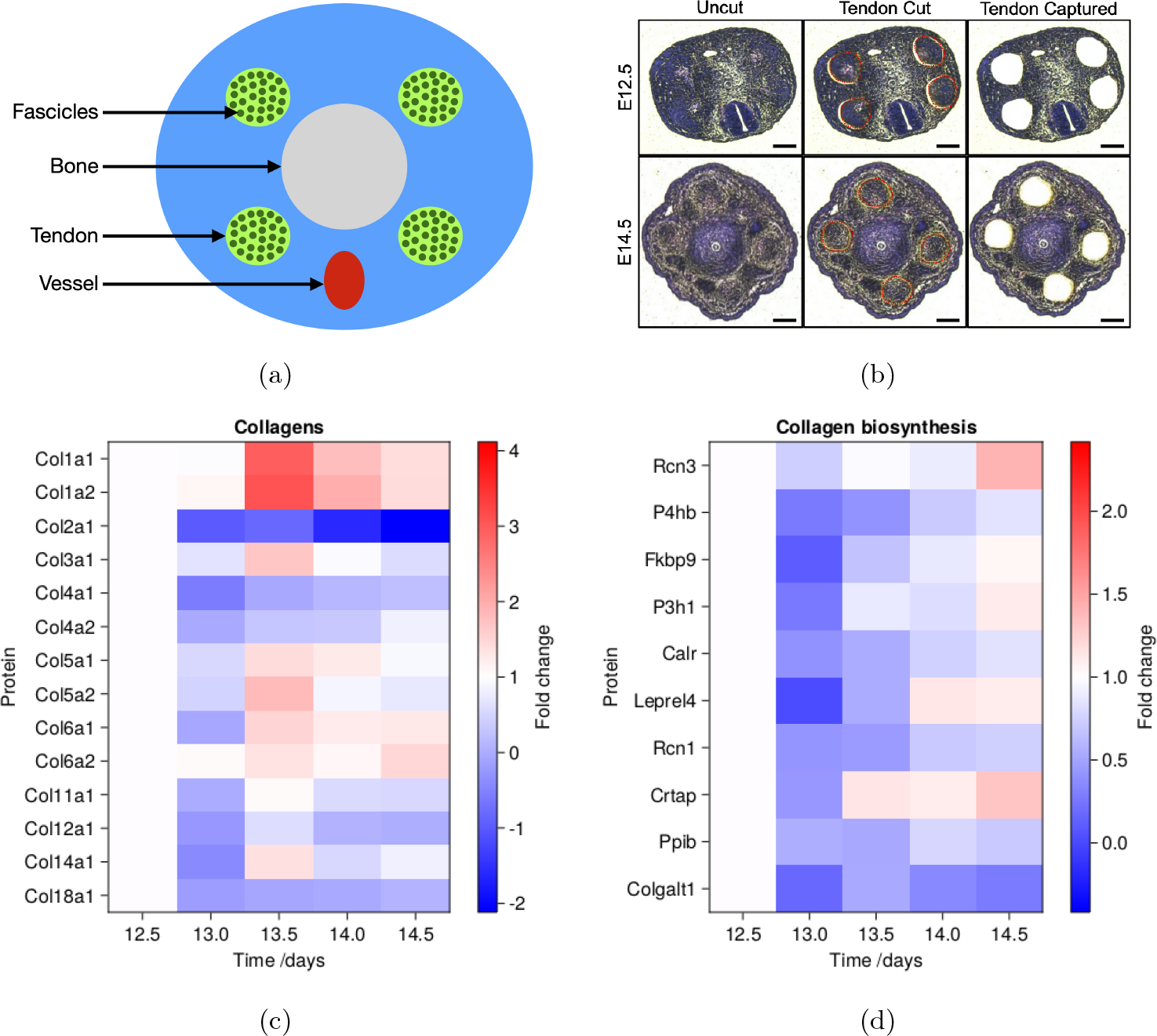
Collagen biosynthesis and collagen proteins increase in tail tendons over time. (a) A cartoon showing the anatomy of a tail with 4 tendons, a central bone and large blood vessel. (b) Shown are representative hematoxylin-&-eosin stained images of embryonic mouse tails at embryonic timepoints E12.5 (upper panels) and E14.5 (lower panels). Tendons were laser-capture microdissected (red dotted circles; middle panels) and captured (right panels) for mass spectrometry preparation. Scale bars 100*µ*m. (c) Plot of log_2_ of fold change in abundance of collagens relative to E12.5. (d) Plot of log_2_ of fold change in abundance of collagen biosynthesis proteins relative to E12.5.

Our approach quantified 2173 proteins across the 5 time points. The proteins were compared to the matrisome project [45] to identify matrix/matrix-associated proteins in our data set; we identified around 20 % of the core matrisome and 5 % of the matrisome-associated proteins. 16 proteins were assigned “collagens”, including protein chains making up collagen-I, −III, −V, −XI, which are central to the formation of the tendon (i.e. collagen fibrils). Further, 30 “ECM glycoproteins”, 7 “proteoglycans”, 18 “ECM regulators”, 18 “ECM affiliated proteins”, and 8 “secreted factors” were also identified and quantified [Supplementary Data 1]. The abundances of matrisome proteins across the time series were then normalised to E12.5; proteins where one or more of the time points do not have a value were excluded. Abundances of detected collagens and proteins involved in collagen biosynthesis are presented as heatmaps [Figure 4c-d]. The full matrisome list can be found in Figure 10 [Appendix G]. A heatmap showing the raw protein intensities and a bar chart showing mean protein abundance across all time points can be found in Figure 11 [Appendix G].

In accordance with EM images showing that collagen bundles start to appear at E13.5, our mass spectrometry analysis shows that proteins that make up collagen-I protomer (i.e. Col1a1 and Col1a2) significantly increase and peak at E13.5 [Figure 4c], but then steadily decrease. Other collagens, specifically those responsible for collagen-I fibril nucleation [46] (i.e. Col5a1, Col5a2) showed a similar trend where they peak at E13.5 and then start to decrease in abundance [Figure 4c]. The reduction in collagen-I molecules may be due to the incorporation of soluble collagen-I protomers (either homotrimeric Col1a1, or heterotrimeric Col1a1/Col1a2) into fibrils, a process that makes collagen harder to be extracted for detection by mass spectrometry [12]. Proteoglycans known to be involved in post-natal tendon development such as decorin, biglycan and lumican were also detected [47, 48]. Interestingly, many proteins involved in collagen biosynthesis peaked at E14.0 and E14.5, despite a drop in soluble collagen-I levels [Figure 4d]. For instance, P3H1, P4Ha1 P4ha2, P4hb, Ppib, Serpinh1, Rcn3 and Crtap are all critical for collagen biosynthesis [49, 50, 51, 52, 53]. This is supportive of the interpretation that collagen-I protomers are still being produced but deposited in an insoluble manner, i.e. within a collagen fibril. Taken together, we interpret the MS data to be supportive of the PFC model, where collagen bundles rapidly appear with a concurrent sharp decrease in free collagen protomers.

## 4 Discussion

The PFC model that we have implemented here represents a substantial simplification of collagen biochemistry. However, subject to a number of assumptions, reviewed below, it provides a computationally tractable tool with a small number of free parameters that allows us to investigate the spatiotemporal development of fibrils in intercellular spaces within embryonic tendon.

A strong assumption in our approach is that fully resolved molecular dynamics in three spatial dimensions, accounting for fine details of protomer structure, will evolve macroscopically with many of the features that are captured by the relatively inexpensive coarse-grained 2D PFC model [Equation (1a), Appendix A]. Pending the outcome of 3D studies, we can evaluate the evidence supporting this proposition. Imaging of fibril organisation in 3D shows a high degree of anisotropy [43], with fibril patterns showing very limited variation along the axis of the tendon. Three factors may be responsible for this. First, protomers are long, thin and reasonably stiff rods, with a 300:1 aspect ratio and a persistence length roughly the same order of magnitude as their length [54, 55]. At high concentrations, they can be expected to aggregate in a nematic liquid-crystalline phase, with rods strongly aligned with their neighbours, as has been observed within intracellular vesicles [23, 56, 57]. Second, the protomers occupy long, slender intercellular channels; this geometric confinement can be expected to promote coherent organisation of the protomer in a liquid crystal phase [24]. Third, mechanical loading along the axis of the nascent tendon can be expected to induce stresses on the protomers that promote coherent alignment and subsequent aggregation [25, 26]. Together, these factors mitigate against the formation of an isotropic gel, as occurs *in vitro* [58], but they support use of a simpler 2D model, addressing evolution in the plane normal to the tail tendon, that captures the aggregation of protomers in intercellular spaces.

This model is agnostic regarding the initial nucleating trigger, as well as active cell-involvement, for fibril formation in the embryonic tendon. Pattern formation in the PFC arises from any initial fluctuation in *ϕ* across the domain, but the equilibrium results appear independent of whether this fluctuation is localised or distributed throughout the domain. We hypothesise that factors such as biochemical interaction with other species such as collagen-V [46], geometric confinement leading to nematic ordering [24], axial strain by newly forming muscles, or reaching some threshold concentration of collagen protomers are possible nucleation candidates. In this current study, we observed the emergence of fibrils at E13.5, which filled the intercellular space by E14. From a biological point of view, factors preventing premature initiation of fibrillogenesis may be several-fold. Collagen-I fibrillogenesis in vitro requires a certain threshold of collagen-I molecules, as well as the presence of nucleators such as collagen-V and collagen-XI [46]. Other factors may include a change in pH in the microenvironment [59], cleavage of the propeptide region [60], or the presence of a yet-unidentified inhibitory molecule against fibrillogenesis. In our data, detection of the known nucleating collagens (collagen-V, collagen-XI) followed the trend of collagen-I, supporting the hypothesis that fibril emergence led to a decrease of free collagen-I protomers. We did not detect the known enzymes for cleavage of the propeptides in procollagen-I (i.e. BMP1, ADAMTS2), thus cannot draw conclusions on whether a synchronised removal of propeptide regions contributed towards the rapid appearance of fibrils. Nonetheless, regardless of what mechanisms are employed by cells to control fibril initiation, the current model does not preclude their existence. Previously, we identified fibripositors in mouse embryonic tendon that supported a cell-directed fibrillogenesis model, where their presence were detected at E14.5 and not at E13.5 [9]. In the present study we observed the emergence of fibrils in E13.5, without the presence of obvious fibripositors; the appearance of fibripositors are only detected from E14 onwards (data not shown). From this we inferred that cell-fibril contacts are less frequent in the initial phase of fibril appearance, suggesting that our alternative phase transition model provides a key mechanism that mediates rapid fibril growth. This is also suggestive of a two-step embryonic tendon development; however, to further test this hypothesis is beyond the scope of this study.

Figure 4 provides experimental evidence of protomer abundance during fibril formation. Here, 3.4% of total proteins detected were matrisome or matrisome-associated, whilst other reports on mouse lungs reported 5.2% [12]. The coverage of matrisome or matrisome-associated proteins is highly variable, which is influenced by sample preparation method, and importantly, tissue type and age. For example, during embryogenesis the majority of the tissue is occupied by cells, a phenomena that is reversed in adult. Regardless, our data demonstrate that there is a steady accumulation of protomeric collagen into the intercellular space from E12.5 up until E13.5. The PFC model simulates the subsequent aggregation of this collagen into fibrils. While the PFC model incorporates randomness in the initial distribution of protomer, it does not directly simulate stochastic effects in the subsequent dynamics. The model predicts a rapid initial phase of aggregation [Figure 2c], in which the free energy and protomer availability drop rapidly, consistent with Figure 4c, followed by a slower phase in which the fibril patterns reorganise slowly to lower-energy states with fewer defects [Figure 2d]. Our simulations illustrate the dynamic nature of fibril patterns, highlighting the fact that EM images represent snapshots of an evolving system. Defects are an intrinsic hallmark of fibril patterns: they are present because perfectly crystalline patterns are incompatible with the irregular shape of domain boundaries, and because of inherent environmental disorder. Our simulations aimed to reproduce domain boundaries accurately (in 2D, at least; we did not account for axial variations in shape that might have a long-range influence). The noise in our model (incorporated through the initial condition, implemented as a Gaussian random field) highlights the importance of interpreting fibril (and defect) patterns in a statistical sense, because individual realisations of the model show variability in the organisation of individual fibrils. While some simulations show encouraging agreement [Figure 3a-d], overall the density of defects predicted by the model underpredicts the density of defects measured in EM images [Figure 3i-k]. Recognising that defect density varies with time, we attribute the difference primarily to 3D effects, to elevated levels of spatial heterogeneity and to possible cross-linking of non-equilibrium fibril patterns, not captured by the model.

The model does not have sufficient degrees of freedom to capture the detailed biochemistry of collagen protomers, including their chirality and specific cross-linkers or molecules controlling fibril diameter (such as lysyl oxidases or decorin) that regulate self-assembly into fibrils [61]. Instead, these effects are aggregated into the free-energy parameter *r* and the initial mean phase field *ϕ*_0_. While there is considerable uncertainty in the true values of these parameters, fibril formation is predicted when the parameters occupy a range of values [Figure 6], indicating the robustness of the proposed mechanism. The most striking effect of parameter variation within this allowable range is the appearance of voids in the fibril pattern (for smaller values of *r* and larger values of *|ϕ*_0_*|*, see also Figure 2e). These reflect the fibril-free voids revealed in many EM images within intercellular spaces [Figure 8]. Additional secretion of protomeric collagen into the intercellular spaces during fibril crystallisation, which may be supposed to correspond to a reduction in mean *ϕ* across the domain, may push the system from an equilibrium with voids towards a more uniform fibril lattice, as illustrated in Figure 6. While 3D effects are likely to be implicated in void formation in some instances, our model provides an additional potential mechanism, namely as a form of so-called localised state [13] in which fibrils can coexist with regions containing unaggregated protomer. Our simulations closely mimic fibril patterns seen in biological samples, and are supported by biochemistry interpreted through our time-series mass spectrometry data; this is indicative of the feasibility of our model occurring in vivo systems, albeit in specific scenarios such as embryonic development.

Defects and voids together have implications for the mechanical properties and function of a tendon. Assembly of fibrils within a highly confined environment can be expected to generate internal (residual) stresses through molecular reorganisation [62, 63] and as a result of geometric frustration [64]. Slow reorganisation of the fibrils, evident in Figure 2b, is a form of plastic deformation that allows some stress relaxation. Likewise, under mechanical loading, the internal structure of a newly formed tendon can be expected to reorganise via migration and interaction of defects, through a form of annealing, allowing adaptation of growing tendon to loaded conditions. While our model describes the rapid initial formation of fibrils via aggregation, it is likely complementary to cell-mediated and cell-controlled fibrillogenesis during this process; in particular, fibripositors [9] are likely to play an important role over long periods, not only in laying down fibrils, but also potentially guiding the orientation of fibril arrays via mechanical loading of individual fibrils [65]. Further work is required to understand how the strongly anisotropic organisation of intercellular spaces ensures coherent patterning of fibrils along the axis of the tendon in 3D, and mechanisms by which fibril patterns evolve as the embryo matures.

In conclusion, simulations with the phase field crystal model, combined with data obtained by laser-capture microdissection and mass spectroscopy revealing a rapid rise and fall in the abundance of specific intercellular collagen protomers, together provide evidence that embryonic tendon fibril formation occurs as a rapid self-assembly process.

## Author contributions

OEJ, KEK conceived the project; CKR, OEJ conceived and developed the mathematical model and image processing; KEK, JC supervised experiments; JC, JAH, YL performed experiments; CL performed mass spectrometry data processing; JC, JAH interpreted biological data; OEJ, CKR, JC, JAH wrote the manuscript.

## A The phase field crystal model

The PFC model in two spatial dimensions describes the evolution of an order-parameter field *ϕ^∗^*(**x***^∗^, t^∗^*) down gradients of free energy; here stars denote dimensional quantities. We adopt a canonical free energy with density 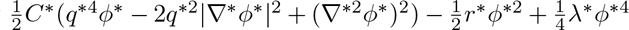, where *C^∗^*, *r^∗^* and *λ^∗^* are positive constants. The corresponding chemical potential density (the first variation of the free energy density, measured in units of energy *k_B_T* per area) is *µ^∗^* = *C^∗^L^∗^*^2^*ϕ^∗^ − r^∗^ϕ^∗^* + *λ^∗^ϕ^∗^*^3^, where *L^∗^ ≡ q^∗^*^2^ + *∇^∗^*^2^. The rate of evolution is regulated by a baseline mobility *α^∗^* that can be expressed as *D^∗^/k_B_T* for some diffusion coefficient *D^∗^*. To express the model in nondimensional terms (minimizing the number of parameters), we measure lengths in terms of 1*/q^∗^*, so that *∇^∗^* = *q^∗^∇*, and time in terms of 1*/*(*α^∗^C^∗^q^∗^*^6^); we define *r* = *r^∗^/*(*C^∗^q^∗^*^4^) (which compares the long-wave-destabilising contribution to the free energy 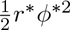 with the term setting the primary lengthscale of the pattern 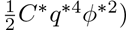) and rescale *ϕ^∗^* using *ϕ^∗^*(**x***^∗^*) = (*C^∗^q^∗^*^4^*/λ^∗^*)^1*/*2^*ϕ*(**x**).

We use the resulting dimensionless phase field *ϕ*(**x**, *t*) to distinguish regions occupied by extracellular collagen protomer, where *ϕ ≈* 1, from regions in which collagen assembles into fibrils, where *ϕ ≈* −1. The free energy is symmetric under the transformation *ϕ → −ϕ*, but we choose a parameter regime where self-assembled collagen forms arrays of spots. The evolution of *ϕ* under the PFC model [13] can then be written as

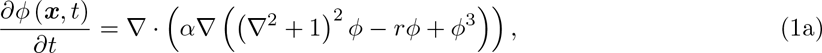

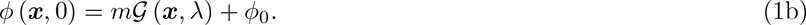

The time-evolution of *ϕ* is thus controlled by the scalar parameter *r*, specifying the destabilising component of the free energy, the mean value of the phase field over the domain *ϕ*_0_, a variance *m*^2^ and a correlation length *λ*. We use the mobility field *α*(**x**) to define accessible (extracellular) and inaccessible (intracellular) regions within solution domains, such that *α* = 1 in the extracellular region and *α* = 0 in the intracellular region. The spatial domain is discussed further in Appendix C. The PFC equation [Equation (1a)] under periodic boundary conditions is conservative, ensuring that the mean *ϕ* over the solution domain is conserved during the dynamics. The initial condition [Equation (1b)] specifies the phase field using a differentiable Gaussian random field *G* (***x***, *λ*) [32] having square-exponential covariance, unit variance, mean 0, and correlation length *λ*. Thus while Equation (1a) is deterministic, disorder enters via the initial condition at a level determined by the amplitude *m*.

The dispersion relation associated with Equation (1a) for small amplitude perturbations around *ϕ* = *ϕ*_0_, having growth rate *σ* and wavenumber *k*, is (with *α* = 1 [13])

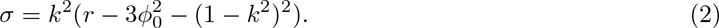

At 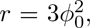, this reveals neutrally stable modes with wavenumbers zero and unity respectively. Elevating *r* above 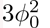 destabilises the long-wave mode, and gives rise to the existence of localised states in which patterned and unpatterned regions may coexist [13]. We hypothesise that this model can thereby explain some features of observations of inter-cellular spaces that are partially filled with fibrils.

Writing the mean nearest-neighbour separation of fibrils in each EM image [Figure 3g] as *ℓ^∗^*, and anticipating from (2) that the (dimensional) wavelength of localised states in the PFC should be *≈* 2*π/q^∗^* (corresponding to *k* = 1), we set 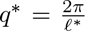. The domain width of an EM image *L^∗^* then appears as the parameter *L* = *q^∗^L^∗^* in simulations.

We focus (mostly) on equilibrium patterns of fibrils and restrict attention to regimes in (*r, ϕ*_0_) parameter space where patterns of spots appear, seeking to understand the credibility of the model in predicting fibril patterns [Figure 6]. We evaluate the robustness of predictions against variations in *r*, *ϕ*_0_ variations in *m* and *λ*.

## B Numerical method

The PFC model [Equation (1)] is 6th-order in space, and therefore challenging to solve computationally over long time intervals. We applied a semi-implicit splitting scheme [33] that introduces a splitting parameter *a* and splits the PDE into two components that together sum to the original PDE. These components are a linear operator *M*, which is treated implicitly, and a nonlinear component *F*, which is treated explicitly, as

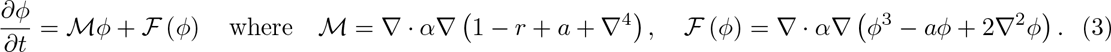

The two terms containing the splitting factor *a* together sum to 0. As recommended in past literature [33], we use a value of *a* = 2.

We construct *M* and *F* from 2nd-order central-difference Laplacians using a Von Neumann neighbourhood 5-point stencil. Due to the spatially varying diffusivity, we must consider the outermost gradient and divergence differential operators (*∇ · α∇*) separately rather than constructing a Laplacian. The *α* field is generated from EM images as described in Appendix C. We developed numerical methods using the Julia programming language [34]. In particular, integration was handled by the Julia DifferentialEquations.jl library [35], using the SplitODEProblem functionality. We used a second-order exponential Runge-Kutta scheme with fixed timestepping, and with a Krylov approximation/operator caching and Krylov subspace of size 50. After discretization, the mean *ϕ* value across the domain was found to be conserved over time to 9 significant figures.

The initial conditions were generated with the Julia GaussianRandomFields.jl package, passing arguments for unit variance and zero mean. Given this Gaussian random field, we set the initial condition to be *ϕ* (***x***, *t* = 0) = 0 for ***x*** within the intra-cellular space (as defined in Appendix C), and *ϕ* (***x***, *t* = 0) = *mG* (***x***, *λ*) + *ϕ*_0_ *− e* for ***x*** within the inter-cellular space. *m* is an amplitude, here set to 0.1, and *e* is the calculated mean of the original Gaussian random field over the inter-cellular domain, such that by subtracting *e* we ensure that the mean phase over the inter-cellular domain is precisely the specified PFC parameter *ϕ*_0_ regardless of individual variation in values produced by each instance of the Gaussian random field.

We rescale the dimensions of an EM image with *x*-dimension length *L^∗^* and horizontal grid spacing *h^∗^* = *L^∗^/M* (for some *M*) to dimensionless length *L* = 2*πL^∗^/l^∗^* (as above) and grid-spacing *h* = (2*πL^∗^*) */* (*l^∗^M*). Simulations were run in batches on the University of Manchester Computational Shared Facility. We use the DrWatson.jl package to manage results produced by testing the parameter space for these simulations [67].

All code is available on GitHub [66].

## C Image processing to define solution domains

To find a domain in which to solve the PFC equation, we started from cross-section EM images of embryonic tail tendon with *x×y* resolution *M ×N* [Figure 5a]. These images were converted to grayscale and binarised to a grid of black and white pixels. Binarised images were segmented to identify structures within the image using the Julia ImageSegmentation.jl package [Figure 5b]. We identified the intercellular and intra-cellular spaces by filtering segments by size to leave only the two largest segments. We converted these segments to a matrix with values of 1 for all pixels in the inter-cellular space and 0 for all other pixels [Figure 5c].

**Figure 5:**
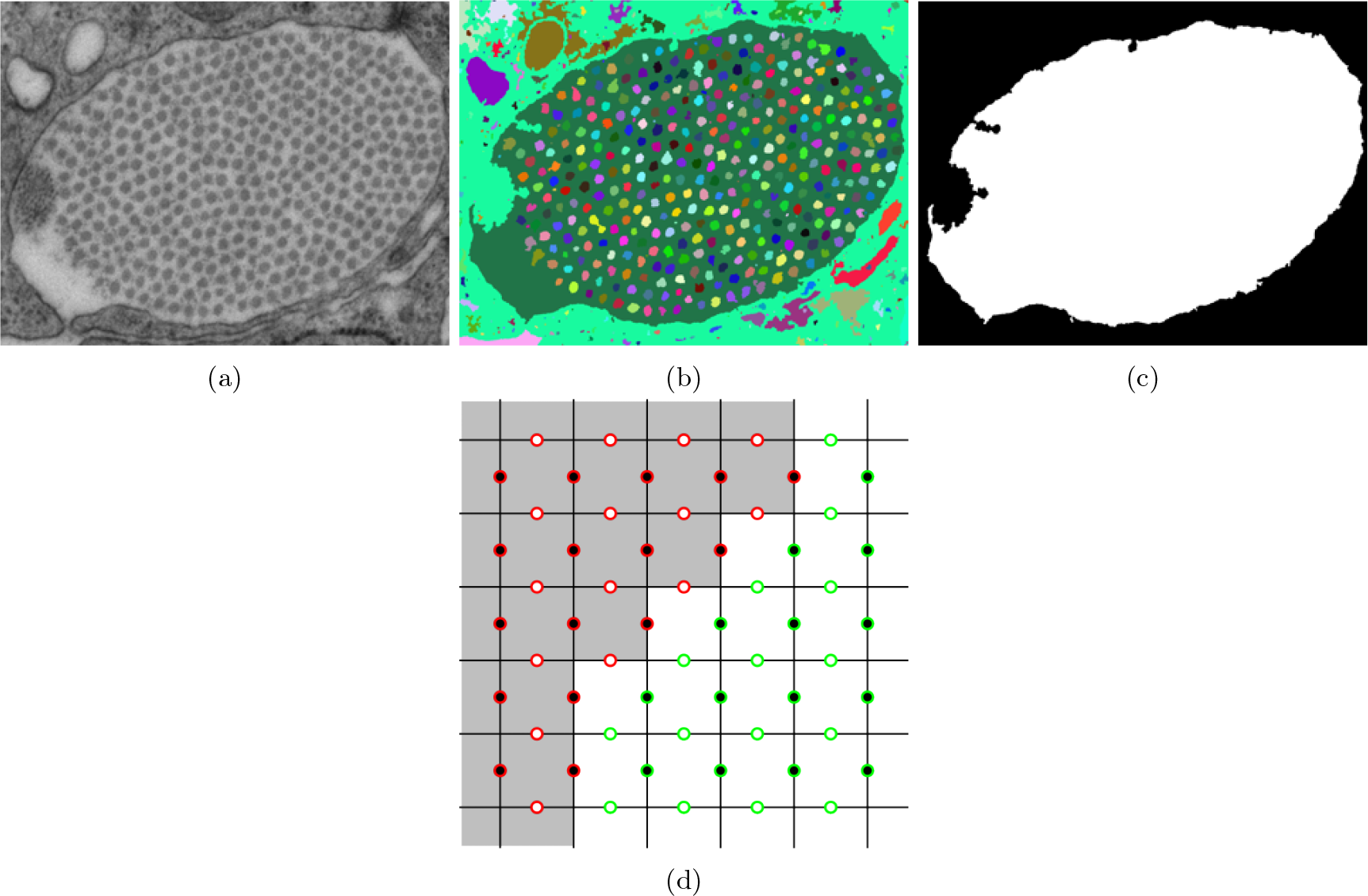
(a-c) Example pipeline showing the progression from an EM image of embryonic tail tendon of *y*-dimension size *N* and *x*-dimension size *M* (a) to a binarised and segmented image (b) and finally a mask of inter-cellular and cellular spaces (c) obtained by pruning smaller segments. (d) Diagram demonstrating how diffusivity matrices are constructed from binarised image mask. Circles represent diffusivity values between pixels in the binarised image. Black filled circles are x-dimension diffusivity values; white-filled circles are y-dimension diffusivity values. Pixel colours represent corresponding black (0) and white (1) components in the binarised image mask. Red circles show where diffusivity values are set to zero; green circles show where diffusivity values are set to 1.

Having created a binarised mask, we set diffusivity values for the PFC solution. We solved Equation (1a) on a periodic domain, but restricted PFC evolution to the inter-cellular space by setting diffusivity to 1 in the inter-cellular space and 0 elsewhere. In practice, this means that diffusivity values were stored as a diagonal matrix *α* of dimension 2*NM ×* 2*NM*, where each diagonal component corresponds to the diffusivity of an edge connecting pixels in the binarised mask. If the value of either pixel is 0 then the diffusivity is 0; if both have a value of 1 then the corresponding diffusivity is 1 [Figure 5d].

## D Parameters

Parameters used in our simulations are outlined in in Table 1. A phase space illustrating the impact of varying *r* and *ϕ*_0_ on the patterns emerging at long times is given in Figure 6. Larger *ϕ*_0_ and smaller *r* promote the appearance of patches in the domain that are not occupied by fibrils. The array of images indicates how the randomness of the initial conditions leads to variability in the patterns of fibrils that emerge, and in the shapes of the fibril-free patches.

**Figure 6:**
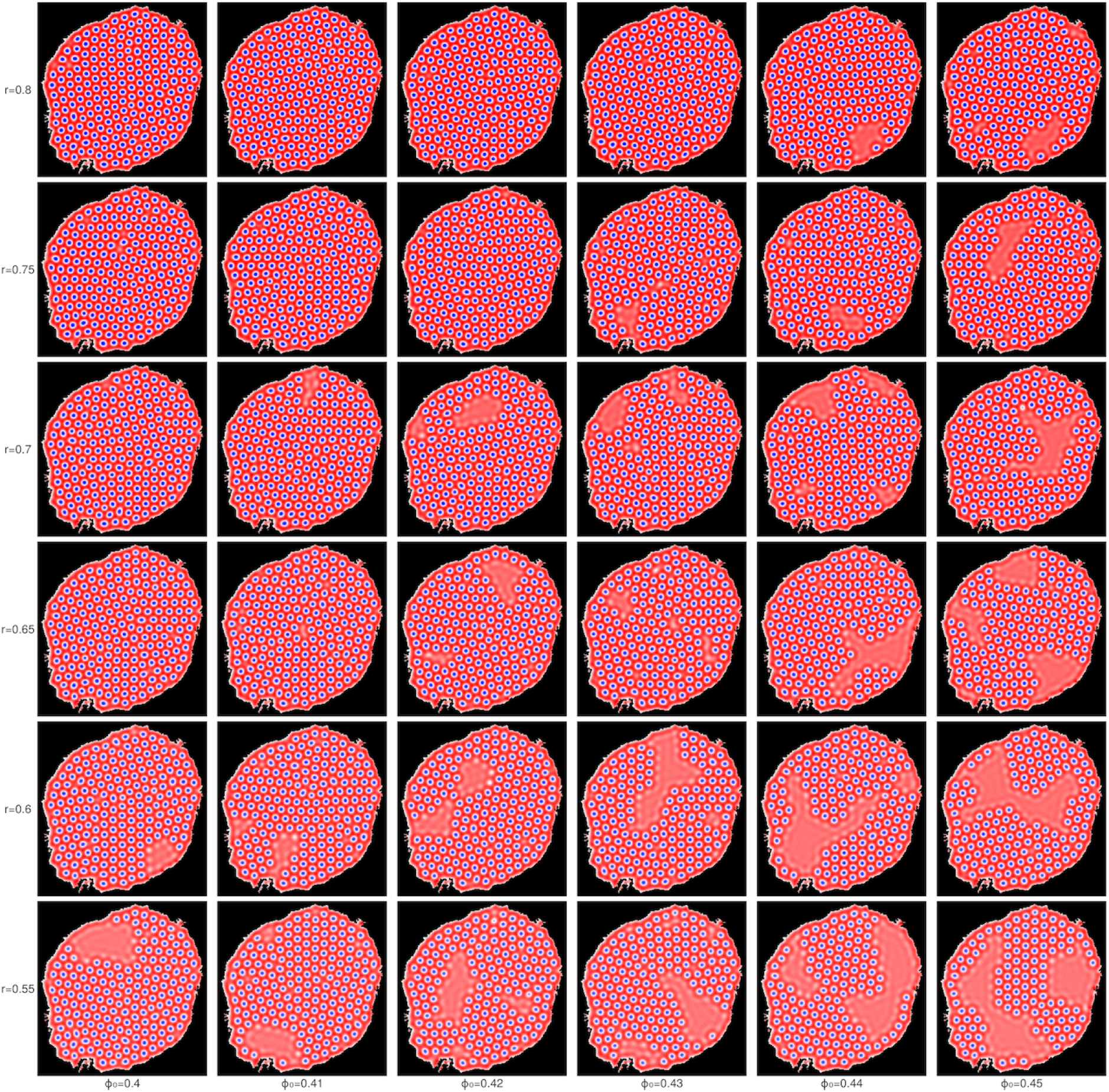
Phase space in *r* and *ϕ*_0_ of PFC solutions in the same solution domain at *t* = 1000. Other parameters are given in Table 1.

**Table 1:** Parameters used to produce results in Section 3.

| Parameter | Value |
| --- | --- |
| $\phi_0$ | 0.4 to 0.45 |
| $r$ | 0.5 to 0.8 |
| $a$ | 2.0 |
| $dt$ | 0.1 |
| $m$ | 0.1 |
| $tMax$ | 1000 |
| $\lambda$ | 10 |

## E Image analysis

To provide a more refined assessment of fibril patterns, we identified the locations of defects in an otherwise hexagonal array of fibrils as follows. Simulation results were processed by binarising simulation results with a threshold of *ϕ* = 0.5. These binarised results were segmented, again using the Julia ImageSegmentation.jl library. Fibril segments were obtained by filtering segments by pixel count, and the centroid position of each fibril was found by calculating the centre of mass of each segment’s pixels. These centroids were analysed in the same manner as fibril locations found manually in EM images.

For both EM images and simulation results, we performed a Delaunay triangulation over the set of centroid locations. We found a concave hull for the set of points using the Julia ConcaveHull.jl package. Fibrils within the concave hull set were considered to be peripheral fibrils and were excluded from subsequent analysis. We also excluded fibrils next to voids inside the lattice by calculating the mean area of internal Voronoi cells and excluding those with area greater than 1.3 times the mean from defect analysis. Using the Delaunay triangulation, we then determined fibril neighbour counts, distances between nearest neighbours, and lattice defect patterns (identifying internal fibrils which do not have 6 neighbours). The defect patterns of the simulations shown in Figure 6 are shown in Figure 7.

**Figure 7:**
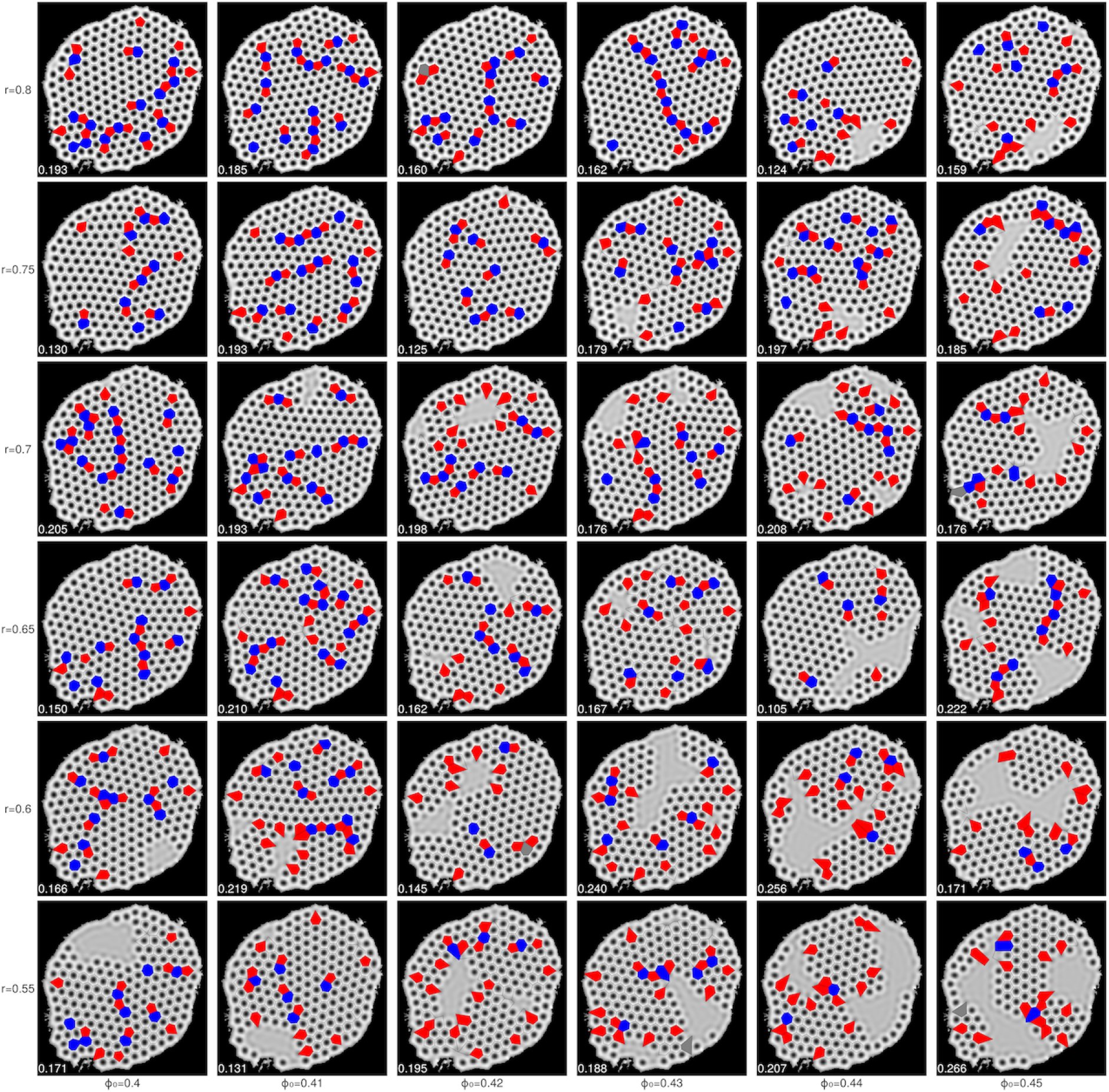
Phase space in *r* and *ϕ*_0_ as in Figure 6 with Voronoi tessellation over cells and defects in neighbour count highlighted. Voronoi cells above a threshold size are excluded to avoid edge effects from empty spaces in the solution. Fibrils with 5 neighbours are coloured red, fibrils with 7 neighbours are coloured blue.

## F Electron micrograph defects

Thirty examples of fibril patterns in EM images were analysed to show defect patterns, using the approach described in Appendix E, as shown in Figure 8. The distributions of neighbour counts and the proportion of fibrils identified as defects are shown in Table 2. Representative simulations run in each of the 30 domains are shown in Figure 9.

**Figure 8:**
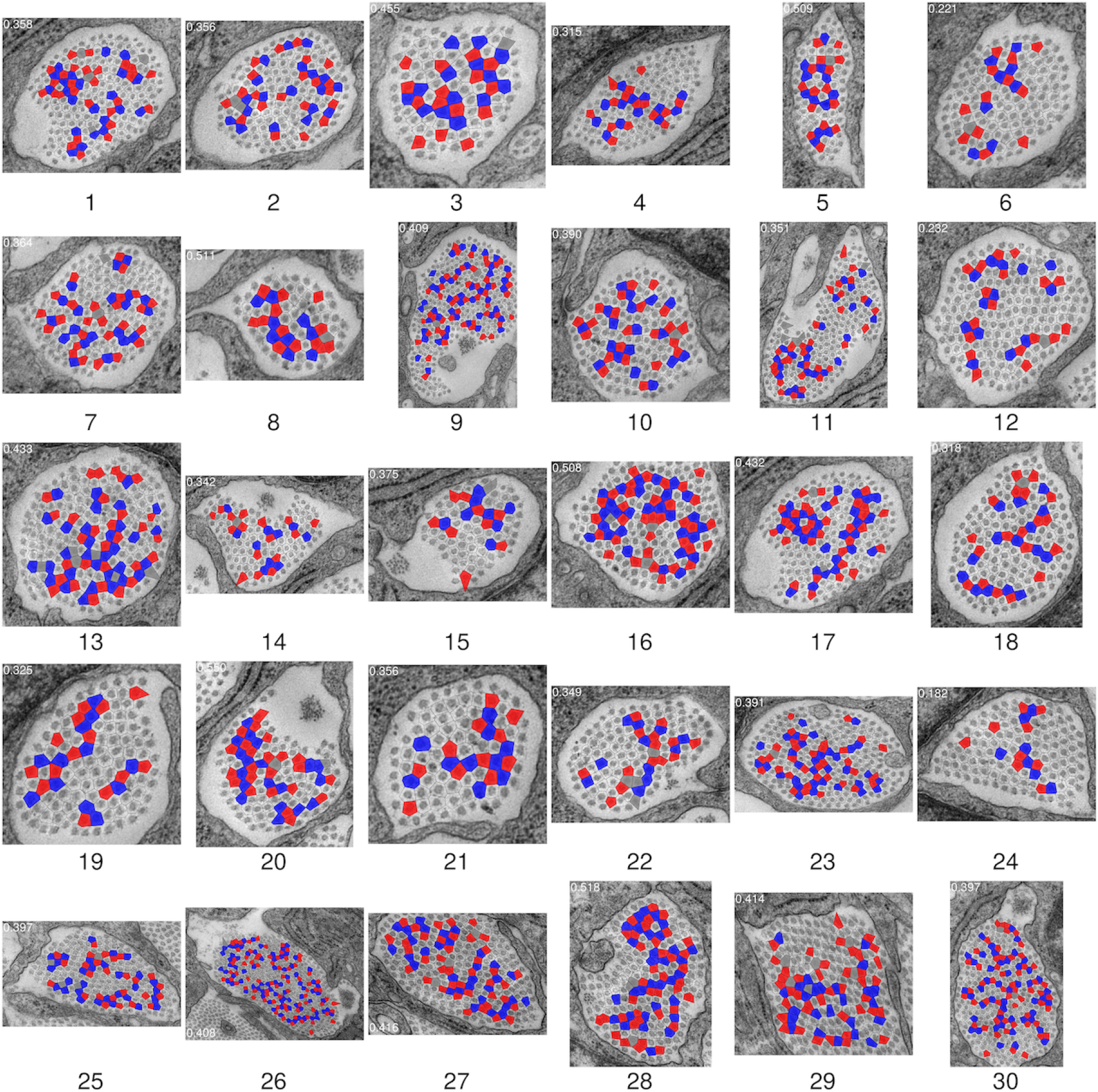
Voronoi tessellation over fibril locations, with Voronoi cells coloured by fibril neighbour count. Fibrils with 6 neighbours are represented by clear cells, 5 neighbours by red cells, and 7 neighbours by blue cells. coloured cells therefore indicate the locations of defects in the fibril lattice. The proportion of cells that are defects is recorded in each panel. The mean defect proportion across 30 examples is 0.386.

**Figure 9:**
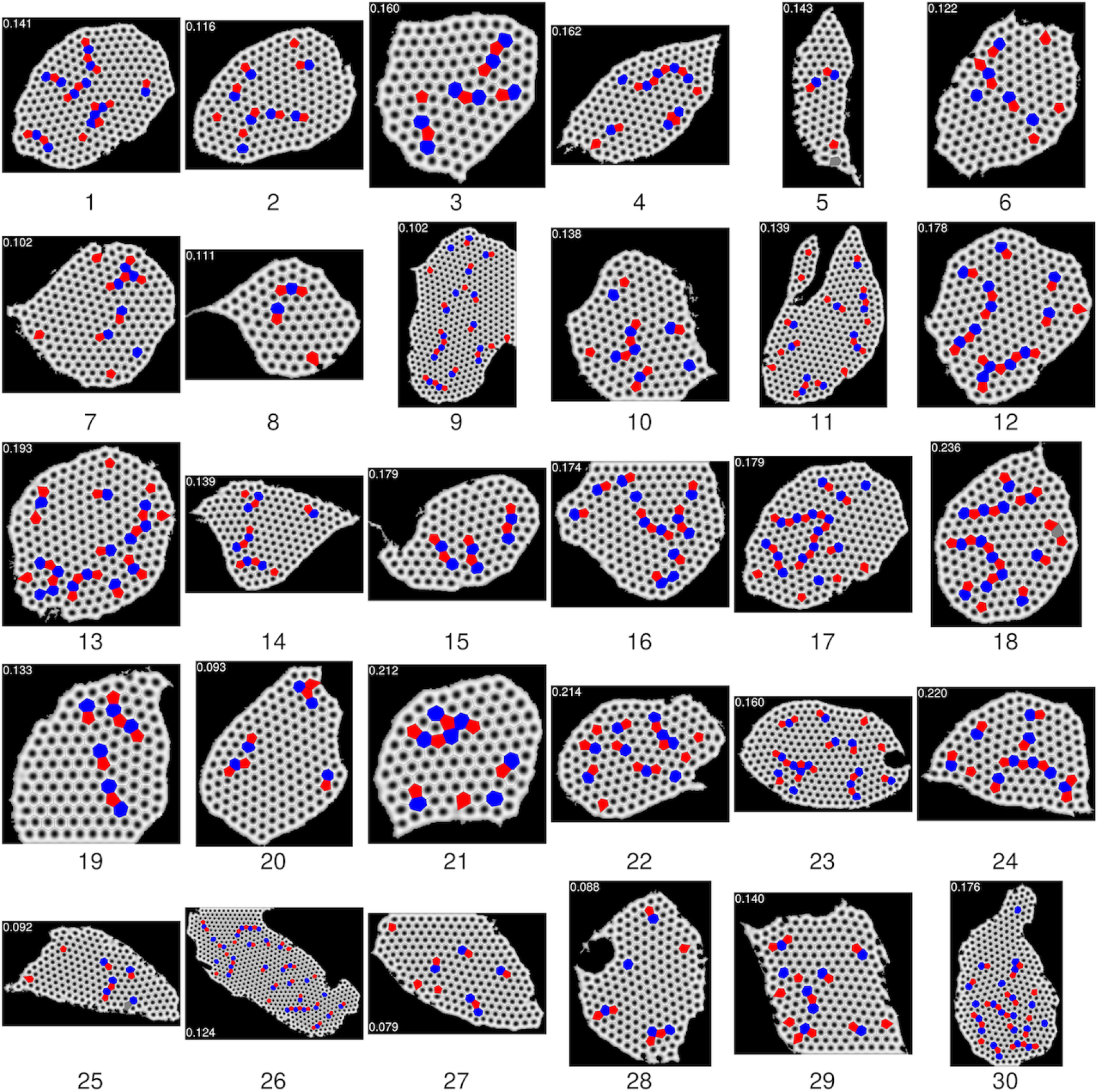
Examples of simulation results for all masks at *r* = 0.8, *ϕ*_0_ = 0.4 and other parameters as in Table 1. The defect proportion in each image is recorded in the corner of the corresponding panel. The mean defect proportion across these runs is 0.148.

**Table 2:** Table specifying neighbour number counts and defect proportions (*d*) for images shown in Figure 8 and corresponding simulations in Figure 9.

| k | EM images |  |  |  |  |  | Simulations |  |  |  |  |  |
| --- | --- | --- | --- | --- | --- | --- | --- | --- | --- | --- | --- | --- |
| | 4 | 5 | 6 | 7 | 8 | $d$ | 4 | 5 | 6 | 7 | 8 | $d$ |
| 1 | 1 | 27 | 95 | 22 | 3 | 0.358 | 0 | 13 | 152 | 12 | 0 | 0.141 |
| 2 | 1 | 21 | 76 | 19 | 1 | 0.356 | 0 | 9 | 122 | 7 | 0 | 0.116 |
| 3 | 1 | 15 | 36 | 14 | 0 | 0.455 | 0 | 6 | 68 | 7 | 0 | 0.160 |
| 4 | 0 | 15 | 61 | 13 | 0 | 0.315 | 0 | 9 | 88 | 8 | 0 | 0.162 |
| 5 | 0 | 15 | 27 | 12 | 1 | 0.509 | 1 | 3 | 36 | 2 | 0 | 0.143 |
| 6 | 0 | 12 | 74 | 8 | 1 | 0.221 | 0 | 7 | 86 | 5 | 0 | 0.122 |
| 7 | 1 | 26 | 84 | 20 | 1 | 0.364 | 0 | 9 | 132 | 6 | 0 | 0.102 |
| 8 | 0 | 13 | 22 | 9 | 1 | 0.511 | 0 | 4 | 48 | 2 | 0 | 0.111 |
| 9 | 2 | 49 | 143 | 46 | 2 | 0.409 | 0 | 18 | 299 | 16 | 0 | 0.102 |
| 10 | 0 | 22 | 64 | 18 | 1 | 0.390 | 0 | 8 | 94 | 7 | 0 | 0.138 |
| 11 | 4 | 30 | 109 | 24 | 1 | 0.351 | 0 | 18 | 179 | 11 | 0 | 0.139 |
| 12 | 0 | 18 | 106 | 13 | 1 | 0.232 | 0 | 14 | 120 | 12 | 0 | 0.178 |
| 13 | 5 | 30 | 89 | 32 | 1 | 0.433 | 0 | 18 | 134 | 14 | 0 | 0.193 |
| 14 | 0 | 16 | 52 | 10 | 1 | 0.342 | 0 | 8 | 93 | 7 | 0 | 0.139 |
| 15 | 1 | 9 | 30 | 8 | 0 | 0.375 | 0 | 6 | 55 | 6 | 0 | 0.179 |
| 16 | 1 | 34 | 63 | 28 | 2 | 0.508 | 0 | 11 | 109 | 12 | 0 | 0.174 |
| 17 | 1 | 32 | 83 | 29 | 1 | 0.432 | 0 | 17 | 147 | 15 | 0 | 0.179 |
| 18 | 0 | 19 | 75 | 15 | 1 | 0.318 | 0 | 16 | 97 | 13 | 1 | 0.236 |
| 19 | 0 | 14 | 54 | 12 | 0 | 0.325 | 0 | 6 | 78 | 6 | 0 | 0.133 |
| 20 | 1 | 23 | 36 | 19 | 1 | 0.550 | 0 | 6 | 107 | 5 | 0 | 0.093 |
| 21 | 0 | 11 | 38 | 10 | 0 | 0.356 | 0 | 7 | 52 | 7 | 0 | 0.212 |
| 22 | 2 | 14 | 54 | 11 | 2 | 0.349 | 0 | 12 | 77 | 9 | 0 | 0.214 |
| 23 | 1 | 35 | 109 | 33 | 1 | 0.391 | 0 | 16 | 157 | 14 | 0 | 0.160 |
| 24 | 0 | 9 | 72 | 7 | 0 | 0.182 | 0 | 11 | 71 | 9 | 0 | 0.220 |
| 25 | 3 | 26 | 91 | 30 | 1 | 0.397 | 1 | 6 | 118 | 5 | 0 | 0.092 |
| 26 | 6 | 77 | 235 | 78 | 1 | 0.408 | 0 | 31 | 425 | 29 | 0 | 0.124 |
| 27 | 0 | 40 | 104 | 31 | 3 | 0.416 | 0 | 7 | 140 | 5 | 0 | 0.079 |
| 28 | 1 | 38 | 68 | 32 | 2 | 0.518 | 0 | 6 | 114 | 5 | 0 | 0.088 |
| 29 | 1 | 41 | 102 | 25 | 5 | 0.414 | 0 | 12 | 123 | 8 | 0 | 0.140 |
| 30 | 0 | 56 | 161 | 48 | 2 | 0.397 | 0 | 20 | 192 | 21 | 0 | 0.176 |

## G Mass spectrometry data processing and analysis

Raw mass spectrometry data were processed using MaxQuant (v.2.0.3.0) [68] against the mouse proteome downloaded from Uniprot (August 2022) [69]. Variable modifications were set as oxidation of methionine, oxidation of proline and protein N-terminal acetylation, with a fixed modifcation as carbamidomethylation of cysteine. The false discovery rate (FDR) was set as 0.01 at both PSM and protein level. Tolerances were set at 20ppm and 4.5ppm for the precursor in both first and main searches, with a 20ppm tolerance set for MS/MS. The option “match between runs” was also selected. Known contaminants were removed. Processed MS data were analysed in R (v.4.1.2) [70] using the R package MSqRob2 (v.1.20) [71].

Peptides were normalised using quantile normalisation and protein roll-up was performed using robust summarisation. Timepoints were compared to the first (E12.5) timepoint, with significantly changing proteins taken at an FDR of 0.05. Heatmaps of relative change in protein abundance over time are shown in Figure 10. Corresponding time-averaged absolute protein abundance across all 5 time points is shown in Figure 11a; a heatmap of absolute protein abundance is shown in Figure 11b.

**Figure 10:**
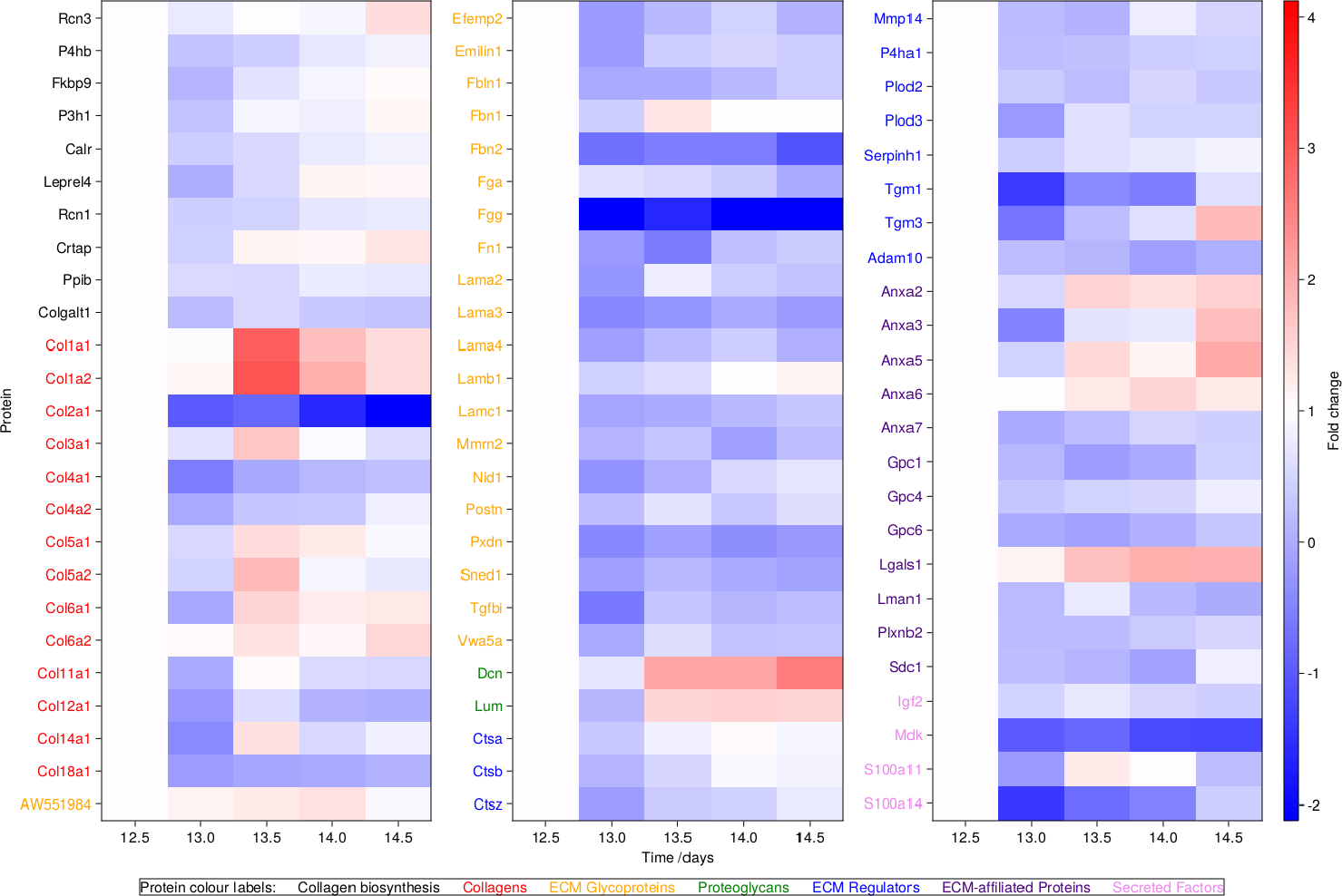
Full heatmap of all relevant protein expression levels over time. Proteins are categorised by function, with different y-axis label colours indicating protein categories. All abundance values are given as log_2_ of fold change, normalised against E12.5, with a white cell indicating no change.

**Figure 11:**
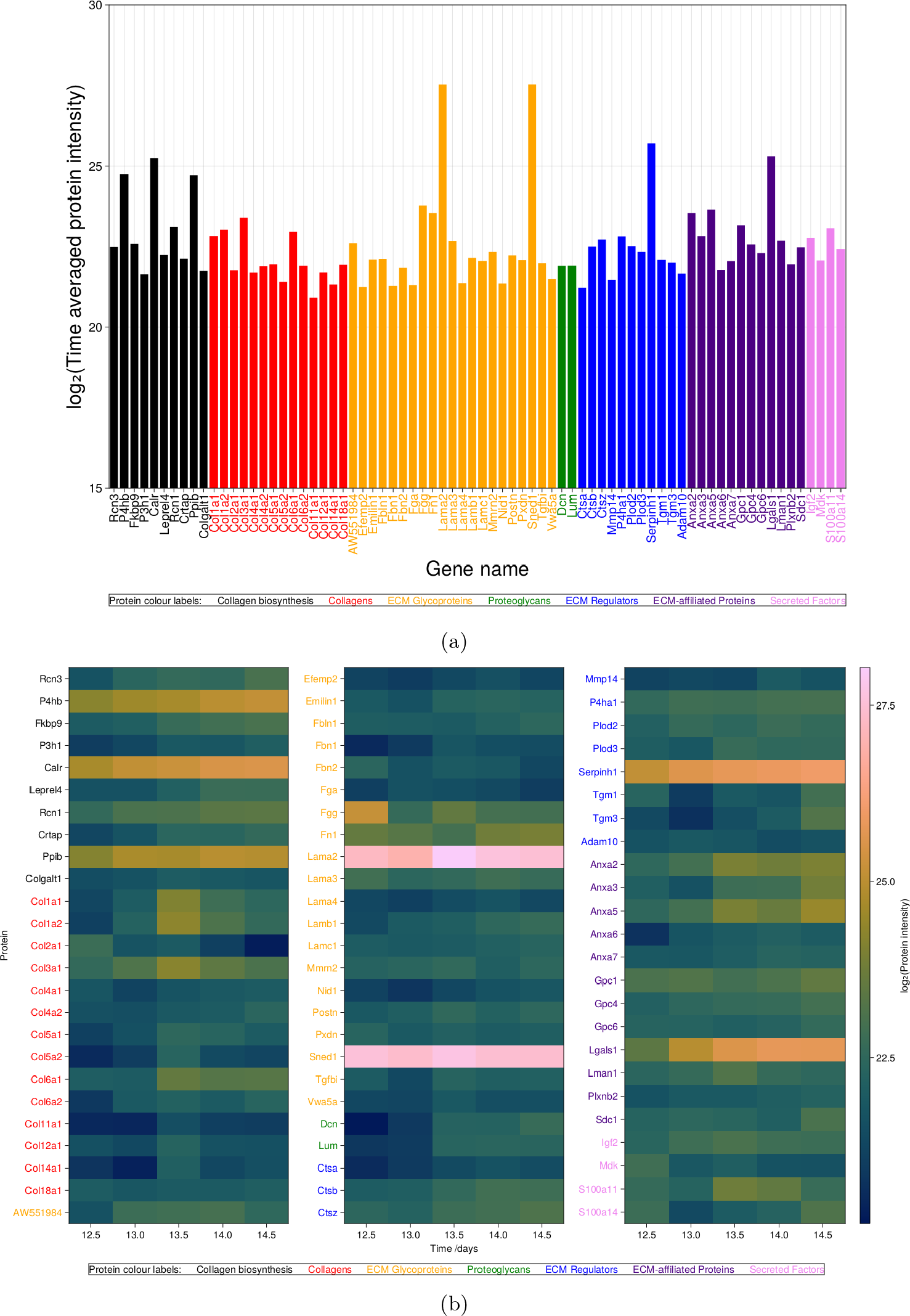
(a) Bar chart showing mean protein abundance across all time points. (b) Heat map showing log_2_ of protein abundance at all timepoints.

## Acknowledgements

The research in the authors’ laboratories is funded by BBSRC (BB/T001984/1), MRC (MR/W016796/1), and The Wellcome Trust (110126/Z/15/Z and 203128/Z/16/Z).

The authors would like to thank Anna Hoyle for additional analysis of the mass spectrometry data, and acknowledge the use of the Computational Shared Facility at The University of Manchester. The proteomic analysis was performed at the Biological Mass Spectrometry Facility.

For the purpose of Open Access, the authors have applied a Creative Commons Attribution (CC BY) licence to any Author Accepted Manuscript version arising.

## Data sharing statement

Mass spec data will be uploaded to ProteomeXchange with unique identifier; meanwhile it will be available upon request to JC. All simualtion and data processing code is available from GitHub [66].

## References

[1] Sylvie Ricard-Blum. The Collagen Family. CSH Perspect. Biol., 3(1), January 2011.

[2] Fransiska Malfait, Marco Castori, Clair A. Francomano, Cecilia Giunta, Tomoki Kosho, and Peter H. Byers. The Ehlers-Danlos syndromes. Nat. Rev. Dis. Primers, 6(1):64, July 2020.

[3] Frank Rauch and Francis H Glorieux. Osteogenesis imperfecta. The Lancet, 363(9418):1377–1385, 2004.

[4] Christopher K. Revell, Oliver E. Jensen, Tom Shearer, Yinhui Lu, David F. Holmes, and Karl E. Kadler. Collagen fibril assembly: New approaches to unanswered questions. Matrix Biol. Plus, 12:100079, December 2021.

[5] Tobias Starborg and Karl E. Kadler. Serial block face-scanning electron microscopy: A tool for studying embryonic development at the cell-matrix interface: SBF-SEM in developmental biology. Birth Defects Res. C, 105(1):9–18, March 2015.

[6] Moses Musiime, Joan Chang, Uwe Hansen, Karl E Kadler, Ćedric Zeltz, and Donald Gullberg. Collagen assembly at the cell surface: dogmas revisited. Cells, 10(3):662, 2021.

[7] Elizabeth G Canty and Karl E Kadler. Procollagen trafficking, processing and fibrillogenesis. J. Cell Sci., 118(7):1341–1353, 2005.

[8] Helen K Graham, David F Holmes, Rod B Watson, and Karl E Kadler. Identification of collagen fibril fusion during vertebrate tendon morphogenesis. The process relies on unipolar fibrils and is regulated by collagen-proteoglycan interaction. J. Mol. Biol., 295(4):891–902, January 2000.

[9] Elizabeth G Canty, Yinhui Lu, Roger S Meadows, Michael K Shaw, David F Holmes, and Karl E Kadler. Coalignment of plasma membrane channels and protrusions (fibripositors) specifies the parallelism of tendon. J. Cell Biol., 165(4):553–563, 2004.

10. Philip W. Anderson. Basic Notions Of Condensed Matter Physics (1st ed.). CRC Press, 1994.

[11] Jeremy A. Herrera, Venkatesh Mallikarjun, Silvia Rosini, Maria Angeles Montero, Craig Lawless, Stacey Warwood, Ronan O’Cualain, David Knight, Martin A. Schwartz, and Joe Swift. Laser capture microdissection coupled mass spectrometry (LCM-MS) for spatially resolved analysis of formalin-fixed and stained human lung tissues. Clin. Proteom., 17(1):24, December 2020.

[12] Nick A. Huizen, Jan N.M. Ijzermans, Peter C. Burgers, and Theo M. Luider. Collagen analysis with mass spectrometry. Mass Spectrom. Rev., 39(4):309–335, July 2020.

[13] Uwe Thiele, Andrew J. Archer, Mark J. Robbins, Hector Gomez, and Edgar Knobloch. Localized states in the conserved Swift-Hohenberg equation with cubic nonlinearity. Phys. Rev. E, 87(4):042915, April 2013.

[14] Nele Moelans, Bart Blanpain, and Patrick Wollants. An introduction to phase-field modeling of microstructure evolution. Calphad, 32(2):268–294, June 2008.

[15] A J Archer, M J Robbins, U Thiele, and E Knobloch. Solidification fronts in supercooled liquids: How rapid fronts can lead to disordered glassy solids. Phys. Rev. E, page 13, 2012.

[16] Long-Qing Chen. Phase-Field Models for Microstructure Evolution. Annu. Rev. Mater. Res., 32(1):113–140, August 2002.

[17] Ĺaszĺo Gŕańasy, Gÿorgy Tegze, Gyula I Tóth, and Tamás Pusztai. Phase-field crystal modelling of crystal nucleation, heteroepitaxy and patterning. Philos. Mag., 91(1):123–149, 2011.

[18] Sven van Teeffelen, Rainer Backofen, Axel Voigt, and Hartmut Löwen. Derivation of the phase-field-crystal model for colloidal solidification. Phys. Rev. E, 79(5):051404, May 2009.

[19] Samuel Cameron, Laurent Kreplak, and Andrew D. Rutenberg. Phase-field collagen fibrils: Coupling chirality and density modulations. Phys. Rev. Res., 2(1):012070, March 2020.

[20] Matthew P. Leighton, Laurent Kreplak, and Andrew D. Rutenberg. Non-equilibrium growth and twist of cross-linked collagen fibrils. Soft Matter, 17(5):1415–1427, 2021.

[21] Heike Emmerich, Hartmut Löwen, Raphael Wittkowski, Thomas Gruhn, Gyula I. Tóth, Gÿorgy Tegze, and Ĺaszĺo Gŕańasy. Phase-field-crystal models for condensed matter dynamics on atomic length and diffusive time scales: An overview. Adv. Phys., 61(6):665–743, December 2012.

[22] Ebrahim Asadi and Mohsen Asle Zaeem. A Review of Quantitative Phase-Field Crystal Modeling of Solid–Liquid Structures. JOM, 67:186–201, 2015.

[23] Marie-Madeleine Giraud-Guille, Laurence Besseau, and Raquel Martin. Liquid crystalline assemblies of collagen in bone and in vitro systems. J. Biomech., 36(10):1571–1579, 2003.

[24] Nima Saeidi, Kathryn P. Karmelek, Jeffrey A. Paten, Ramin Zareian, Elaine DiMasi, and Jeffrey W. Ruberti. Molecular crowding of collagen: A pathway to produce highly-organized collagenous structures. Biomaterials, 33(30):7366–7374, October 2012.

[25] AS Abhilash, Brendon M Baker, Britta Trappmann, Christopher S Chen, and Vivek B Shenoy. Remodeling of fibrous extracellular matrices by contractile cells: predictions from discrete fiber network simulations. Biophys. J., 107(8):1829–1840, 2014.

[26] Georgios Grekas, Maria Proestaki, Phoebus Rosakis, Jacob Notbohm, Charalambos Makridakis, and Guruswami Ravichandran. Cells exploit a phase transition to mechanically remodel the fibrous extracellular matrix. J. Roy. Soc. Interface, 18(175):20200823, 2021.

[27] Jiexiang Lin, Yanping Shi, Yutao Men, Xin Wang, Jinduo Ye, and Chunqiu Zhang. Mechanical Roles in Formation of Oriented Collagen Fibers. Tissue Eng. Part B, 26(2):116–128, April 2020.

[28] Jesus Bueno, Ilya Starodumov, Hector Gomez, Peter Galenko, and Dmitri Alexandrov. Three dimensional structures predicted by the modified phase field crystal equation. Comp. Mater. Sci., 111:310–312, January 2016.

[29] R Prieler, J Hubert, D Li, B Verleye, R Haberkern, and H Emmerich. An anisotropic phase-field crystal model for heterogeneous nucleation of ellipsoidal colloids. J. Phys. Condens. Mat., 21(46):464110, November 2009.

[30] Mohsen Asle Zaeem, Sasan Nouranian, and Mark F Horstemeyer. Simulation of Polymer Crystal Growth with Various Morphologies Using a Phase-Field Model. In Proceedings of the 2012 AIChE Annual Meeting. American Institute of Chemical Engineers, oct 2012.

[31] Andrew D Rutenberg, Aidan I Brown, and Laurent Kreplak. Uniform spatial distribution of collagen fibril radii within tendon implies local activation of pC-collagen at individual fibrils. Phys. Biol., 13(4):046008, August 2016.

32. Gabriel J Lord, Catherine E Powell, and Tony Shardlow. An introduction to computational stochastic PDEs, volume 50. Cambridge University Press, 2014.

[33] Karl Glasner and Saulo Orizaga. Improving the accuracy of convexity splitting methods for gradient flow equations. J. Comput. Phys., 315:52–64, June 2016.

[34] Jeff Bezanson, Alan Edelman, Stefan Karpinski, and Viral B. Shah. Julia: A Fresh Approach to Numerical Computing. SIAM Rev., 59(1):65–98, January 2017.

[35] Christopher Rackauckas and Qing Nie. DifferentialEquations.jl – a performant and feature-rich ecosystem for solving differential equations in Julia. J. Open Res. Software, 5(1):15, 2017.

[36] Franz Aurenhammer. Voronoi diagrams—a survey of a fundamental geometric data structure. ACM Comput. Surv., 23(3):345–405, 1991.

[37] Francisco Ĺopez Jiménez, Norbert Stoop, Romain Lagrange, Jörn Dunkel, and Pedro M. Reis. Curvature-Controlled Defect Localization in Elastic Surface Crystals. Phys. Rev. Lett., 116(10):104301, March 2016.

[38] Jeremy A. Herrera, Lewis Dingle, M. Angeles Montero, Rajamiyer V. Venkateswaran, John F. Blaikley, Craig Lawless, and Martin A. Schwartz. The UIP/IPF fibroblastic focus is a collagen biosynthesis factory embedded in a distinct extracellular matrix. JCI Insight, 7(16):e156115, August 2022.

[39] Ross T. Kohrs, Chunfeng Zhao, Yu-Long Sun, Gregory D. Jay, Ling Zhang, Matthew L. Warman, Kai-Nan An, and Peter C. Amadio. Tendon fascicle gliding in wild type, heterozygous, and lubricin knockout mice: Lubricin affects tendon fascicle gliding. J. Orthop. Res., 29(3):384–389, March 2011.

[40] Jeremy A. Herrera, Lewis A. Dingle, M. Angeles Montero, Rajamiyer V. Venkateswaran, John F. Blaikley, Felice Granato, Stella Pearson, Craig Lawless, and David J. Thornton. The uip honeycomb airway cells are the site of mucin biogenesis with deranged cilia. bioRxiv, 2022.

[41] William Irvine, Vincenzo Vitelli, and Paul M Chaikin. Pleats in crystals on curved surfaces. Nature, 468(7326):947–951, 2010.

[42] Simon Danisch and Julius Krumbiegel. Makie.jl: Flexible high-performance data visualization for julia. J. Open Source Softw., 6(65):3349, 2021.

[43] Tobias Starborg, Nicholas S Kalson, Yinhui Lu, Aleksandr Mironov, Timothy F Cootes, David F Holmes, and Karl E Kadler. Using transmission electron microscopy and 3View to determine collagen fibril size and three-dimensional organization. Nat. Protoc., 8(7):1433–1448, July 2013.

[44] Seyed Mohammad Siadat, Alexandra A. Silverman, Charles A. DiMarzio, and Jeffrey W. Ruberti. Measuring collagen fibril diameter with differential interference contrast microscopy. J. Struct. Biol., 213(1):107697, 2021.

[45] Alexandra Naba, Karl R. Clauser, Sebastian Hoersch, Hui Liu, Steven A. Carr, and Richard O. Hynes. The Matrisome: In Silico Definition and In Vivo Characterization by Proteomics of Normal and Tumor Extracellular Matrices *. Mol. Cell. Proteomics, 11(4), April 2012.

[46] Richard J. Wenstrup, Simone M. Smith, Jane B. Florer, Guiyun Zhang, David P. Beason, Robert E. Seegmiller, Louis J. Soslowsky, and David E. Birk. Regulation of Collagen Fibril Nucleation and Initial Fibril Assembly Involves Coordinate Interactions with Collagens V and XI in Developing Tendon. J. Biol. Chem., 286(23):20455–20465, June 2011.

[47] Guiyun Zhang, Yoichi Ezura, Inna Chervoneva, Paul S. Robinson, David P. Beason, Ehren T. Carine, Louis J. Soslowsky, Renato V. Iozzo, and David E. Birk. Decorin regulates assembly of collagen fibrils and acquisition of biomechanical properties during tendon development. J. Cell. Biochem., 98(6):1436–1449, 2006.

[48] Yoichi Ezura, Shukti Chakravarti, Åke Oldberg, Inna Chervoneva, and David E. Birk. Differential Expression of Lumican and Fibromodulin Regulate Collagen Fibrillogenesis in Developing Mouse Tendons. J. Cell Biol., 151(4):779–788, November 2000.

[49] Shinya Ito and Kazuhiro Nagata. Biology of Hsp47 (Serpin H1), a collagen-specific molecular chaperone. Sem. Cell Dev. Biol., 62:142–151, February 2017.

[50] Na Rae Park, Snehal S. Shetye, Igor Bogush, Douglas R. Keene, Sara Tufa, David M. Hudson, Marilyn Archer, Ling Qin, Louis J. Soslowsky, Nathaniel A. Dyment, and Kyu Sang Joeng. Reticulocalbin 3 is involved in postnatal tendon development by regulating collagen fibrillogenesis and cellular maturation. Sci. Rep., 11(1):10868, December 2021.

[51] Aileen M. Barnes, Weizhong Chang, Roy Morello, Wayne A. Cabral, MaryAnn Weis, David R. Eyre, Sergey Leikin, Elena Makareeva, Natalia Kuznetsova, Thomas E. Uveges, Aarthi Ashok, Armando W. Flor, John J. Mulvihill, Patrick L. Wilson, Usha T. Sundaram, Brendan Lee, and Joan C. Marini. Deficiency of Cartilage-Associated Protein in Recessive Lethal Osteogenesis Imperfecta. New Eng. J. Med., 355(26):2757–2764, December 2006.

[52] Taina Pihlajaniemi, Raili Myllylä, and Kari I. Kivirikko. Prolyl 4-hydroxylase and its role in collagen synthesis. J. Hepatol., 13:S2–S7, January 1991.

[53] Belinda Schegg, Andreas J. Hülsmeier, Christoph Rutschmann, Charlotte Maag, and Thierry Hennet. Core Glycosylation of Collagen Is Initiated by Two *β*(1-O)Galactosyltransferases. Mol. Cell. Biol., 29(4):943–952, February 2009.

[54] Nagmeh Rezaei, Aaron Lyons, and Nancy R Forde. Environmentally controlled curvature of single collagen proteins. Biophys. J., 115(8):1457–1469, 2018.

[55] Karen Claire and R. Pecora. Translational and Rotational Dynamics of Collagen in Dilute Solution. J. Phys. Chem. B, 101(5):746–753, January 1997.

[56] Emmanuel Belamie, Gervaise Mosser, Fŕedéric Gobeaux, and Marie-Madeleine Giraud-Guille. Possible transient liquid crystal phase during the laying out of connective tissues: *α*-chitin and collagen as models. J. Phys.: Cond. Matt., 18(13):S115, 2006.

[57] Marie Madeleine Giraud-Guille, Gervaise Mosser, and Emmanuel Belamie. Liquid crystallinity in collagen systems in vitro and in vivo. Curr. Op. Coll. Interface Sci., 13(4):303–313, 2008.

[58] Gabor Forgacs, Stuart A Newman, Bernhard Hinner, Christian W Maier, and Erich Sackmann. Assembly of collagen matrices as a phase transition revealed by structural and rheologic studies. Biophys. J., 84(2):1272–1280, 2003.

[59] Jonathan BL Bard and John A Chapman. Polymorphism in collagen fibrils precipitated at low p h. Nature, 219(5160):1279–1280, 1968.

[60] Karl E Kadler, Yoshio Hojima, and DJ Prockop. Assembly of collagen fibrils de novo by cleavage of the type i pc-collagen with procollagen c-proteinase. assay of critical concentration demonstrates that collagen self-assembly is a classical example of an entropy-driven process. J. Biol. Chem., 262(32):15696–15701, 1987.

[61] David JS Hulmes. Building collagen molecules, fibrils, and suprafibrillar structures. J. Struct. Biol., 137(1-2):2–10, 2002.

[62] Marjan Shayegan and Nancy R Forde. Microrheological characterization of collagen systems: from molecular solutions to fibrillar gels. PloS one, 8(8):e70590, 2013.

[63] Max Renner-Rao, Franziska Jehle, Tobias Priemel, Emilie Duthoo, Peter Fratzl, Luca Bertinetti, and Matthew J Harrington. Mussels fabricate porous glues via multiphase liquid–liquid phase separation of multiprotein condensates. ACS nano, 16(12):20877–20890, 2022.

[64] Gregory M Grason. Perspective: Geometrically frustrated assemblies. J. Chem. Phys., 145(11):110901, 2016.

[65] Jiexiang Lin, Yanping Shi, Yutao Men, Xin Wang, Jinduo Ye, and Chunqiu Zhang. Mechanical roles in formation of oriented collagen fibers. Tissue Eng. B: Reviews, 26(2):116–128, 2020.

[66] Christopher K. Revell. PhaseFieldCrystal.jl. https://github.com/chris-revell/ PhaseFieldCrystal, 2023.

[67] George Datseris, Jonas Isensee, Sebastian Pech, and Tamás Gál. DrWatson: The perfect sidekick for your scientific inquiries. J. Open Source Software, 5(54):2673, October 2020.

[68] Stefka Tyanova, Tikira Temu, and Juergen Cox. The maxquant computational platform for mass spectrometry-based shotgun proteomics. Nat. Protoc., 11(12):2301–2319, 2016.

[69] The UniProt Consortium. UniProt: the universal protein knowledgebase in 2021. Nucleic Acids Res., 49(D1):D480–D489, 11 2020.

[70] R R Core Team, et al. R: A language and environment for statistical computing. 2013.

[71] Ludger JE Goeminne, Kris Gevaert, and Lieven Clement. Experimental design and data-analysis in label-free quantitative lc/ms proteomics: A tutorial with msqrob. J. Proteomics, 171:23–36, 2018.

